# Plethora of QTLs found in *Arabidopsis thaliana* reveals complexity of genetic variation for photosynthesis in dynamic light conditions

**DOI:** 10.1101/2022.11.13.516256

**Authors:** Tom P.J.M. Theeuwen, Louise L. Logie, Sanne Put, Hedayat Bagheri, Konrad Łosiński, Justine Drouault, Pádraic J. Flood, Corrie Hanhart, Frank F.M. Becker, Raúl Wijfjes, David Hall, David M. Kramer, Jeremy Harbinson, Mark G.M. Aarts

## Abstract

The environments in which plant species evolved are now generally understood to be dynamic rather than static. Photosynthesis has to operate within these dynamic environments, such as sudden changes to light intensities. Plants have evolved photoprotection mechanisms that prevent damage caused by sudden changes to high light intensities. The extent of genetic variation within plants species to deal with these dynamic light conditions remains largely unexplored. Here we show that one accession of *A. thaliana* has a more efficient photoprotection mechanism in dynamic light conditions, compared to six other accessions. The construction of a doubled haploid population and subsequent phenotyping in a dynamically controlled high-throughput system reveals up to 15 QTLs for photoprotection. Identifying the causal gene underlying one of the major QTLs shows that an allelic variant of *cpFtsY* results in more efficient photoprotection under high and fluctuating light intensities. Further analyses reveal this allelic variant to be overprotecting, reducing biomass in a range of dynamic environmental conditions. This suggests that within nature, adaptation can occur to more stressful environments and that revealing the causal genes and mechanisms can help improve the general understanding of photosynthetic functioning. The other QTLs possess different photosynthetic properties, and thus together they show how there is ample intraspecific genetic variation for photosynthetic functioning in dynamic environments. With photosynthesis being one of the last unimproved components of crop yield, this amount of genetic variation for photosynthesis forms excellent input for breeding approaches. In these breeding approaches, the interactions with the environmental conditions should however be precisely assessed. Doing so correctly, allows us to tap into nature’s solution to challenging environmental conditions.

## Introduction

In natural habitats, plants are at complete mercy of the dynamic properties of the environmental conditions, which are highly dynamic even in agricultural systems. Especially photosynthesis is highly responsive to environmental conditions (Anderson *et al*., 1995). Fluctuating light conditions determine the overall functioning of photosynthesis in crops to a large extent. Clouds passing by cause sudden drops in light intensity, while wind inside canopies causes sudden spikes in light intensity due to leaf movements (Kaiser *et al*., 2018; Durand *et al*., 2021). Photosynthesis is able to work efficiently at many different light intensities, yet adaptation to sudden changes to light intensity takes time. Under high light conditions, to avoid too much light reaching the photosystems, plants can dissipate this excess energy as heat in a process called non-photochemical quenching (NPQ). NPQ in higher plants can grossly be divided into two components, the so called rapidly relaxing (*q*_E_) and slowly relaxing (*q*_I_) components (Müller *et al*., 2001). The *q*_E_ component is the result of energy-dependent quenching, while the *q*_I_ component is the result of photoinhibition, the xanthophyll cycle, state transitions and chloroplast movements (Cruz *et al*., 2016). The fast response of NPQ to a change in light intensity relies on conformational changes in the light harvesting complex being disentangled from the reaction centres. While NPQ is a very dynamic process the relaxation of NPQ can be slow in high to low light transitions. This results in too much energy being dissipated as heat that could otherwise be used for photosynthesis. Modelling these losses in crops shows that this can result in a drop of CO_2_ fixation of up to 30% (Zhu *et al*., 2004).

Since the core photosynthetic machinery is rather conserved, little functional genetic variation is available, that can be used to improve photosynthetic functioning. As a result most improvement studies have focused on overexpressing, mutating or inserting genes known to be involved in the core mechanisms known to be involved in photosynthesis (Ort *et al*., 2015). This is also true for improvements in NPQ dynamics, where overexpression of genes has been shown to bring about conformational changes of the antenna complexes (Johnson *et al*., 2008). Accelerating the NPQ relaxation in tobacco and soybean has been shown to result in 15% and 30% higher yields respectively (Kromdijk *et al*., 2016; De Souza *et al*., 2022). The same approach in *Arabidopsis thaliana* and potato did not result in accelerated NPQ relaxation or changes in yields, showing that it is not a one-size-fits-all solution (Garcia-Molina and Leister, 2020; Lehretz *et al*., 2022). Despite the relative absence of natural genetic variation in the core photosynthetic machinery, there is ample phenotypic variation for photosynthesis, implying there is standing genetic variation outside the core machinery. This is also true for NPQ dynamics, as quantitative trait loci (QTLs) are identified for NPQ in regions of the genome that do not include any of the known NPQ related genes (Poormohammad Kiani *et al*., 2008; Jung and Niyogi, 2009; Wang *et al*., 2017; Oakley *et al*., 2018*a*; Rungrat *et al*., 2019; Goto *et al*., 2021). Unfortunately, in hardly any of these cases the causal genes have been revealed, even though identifying the genes underlying QTLs opens up novel targets for improving the dynamic responses of NPQ, as well as forming an opportunity to expand the physiological understanding of NPQ (Theeuwen *et al*., 2022*b*).

Whether the identified QTLs in any of the previous studies captured all the genetic variation for NPQ present within the population is difficult to assess. To reveal all genetic variation within a population, the choice of mapping populations and the high-throughput phenotyping systems are critical. Currently, genome wide association studies (GWAS) are the preferred approach to reveal novel QTLs for a trait of interest. GWAS are often considered successful when they reveal one or two QTLs. However, due to the need for correction for multiple testing, false positives are difficult to distinguish from true positives. GWAS are also known for their poor statistical power to detect the effects of rare alleles. The difficulty in revealing true positives and poor statistical power results in only a fraction of the QTLs present within the population being revealed (Theeuwen *et al*., 2022*b*). Alternatively bi- or multiparental mapping populations can be used. These populations generally segregate for less genetic variants, in comparison to populations used in GWAS. In bi- or multiparental mapping populations any allele segregates at roughly equal ratios, increasing statistical power to detect QTLs. This results in more QTLs being detected and thus generates a better overview of how much natural genetic variation for NPQ is present within the population. Revealing natural genetic variation for NPQ also depends on the high-throughput phenotyping method used. To follow NPQ as it dynamically responds to changes in environment, continuous phenotyping of the entire mapping population is required (Murchie *et al*., 2018; Bezouw *et al*., 2019). Combining the benefits of revealing more QTLs via a bi- or multiparental populations with controlled dynamic environment phenotyping facilities allows us to reveal how natural genetic variation for dynamic photosynthesis manifests itself.

The *A. thaliana* accession Ely is known to possess a mutation in the chloroplast encoded *PsbA* gene (El-Lithy *et al*., 2005), but here we discovered the nuclear counterpart of this accession to have a more efficient NPQ mechanism. To reveal the underlying nuclear genetic variation for the observed differences in NPQ, a large doubled haploid population between Ely and the common reference genotype Col was constructed. QTL mapping was done with phenotypes collected in a dynamic environment high-throughput phenotyping system. Subsequent candidate gene validation of two QTLs revealed novel physiological insights in dynamic photosynthetic properties.

## Results

### Revealing an efficient NPQ mechanism

The *A. thaliana* accession Ely is known to carry a *PsbA* mutation in the chloroplast genome, but the large phenotypic effect of the *PsbA* mutation masks possible variation for photosynthesis encoded in the nuclear genome. Separating the phenotypic response associated with the mitochondrial and chloroplast genomes (i.e. the plasmotype) from the phenotypic response associated with the nuclear genome (i.e. the nucleotype) is difficult. Recently, a quick and efficient method to make cybrids was developed (Flood & Theeuwen *et al*., 2020). Cybrids are novel combinations between the nucleotype of one accession with the plasmotype of another accession. A previously constructed cybrid panel of seven *A. thaliana* accessions, including the Ely accession, enables the separation of the nucleotype effect of Ely from the *PsbA* mutation. Flood & Theeuwen *et al*., 2020 grew the cybrid panel in a high-throughput phenotyping system able to simulate dynamic environmental conditions, in both stable and fluctuating light conditions (Figure 1A). Here, we reanalysed the phenotypic data and find that the cybrids with the Ely *PsbA* mutation differ significantly from cybrids with wild-type *PsbA* allele in both stable and dynamic light conditions for all measured photosynthetic parameters (*Φ*_PSII_, *Φ*_NPQ_, *Φ*_NO_, NPQ, *q*_E_ and *q*_I_) (Figure 1B and Supplementary Figure 1). The biggest reduction in NPQ between the cybrids with the *PsbA* mutation and without occurred during highly fluctuating light conditions (Figures 1B). However, the NPQ of Ely^Ely^ (noted as Nucleotype^Plasmotype^) was found to be comparable to Col^Col^, even though the Ely^Ely^ genotype has the plasmotypic *PsbA* mutation (Figure 1C). This compensation is caused by the Ely nucleotype, which results in up to 28.6% higher NPQ in comparison to the Col nucleotype (Figure 1C). This means that we revealed the Ely nucleotype to have a different capacity to do NPQ compared to the other nucleotypes.

**Figure 1.**
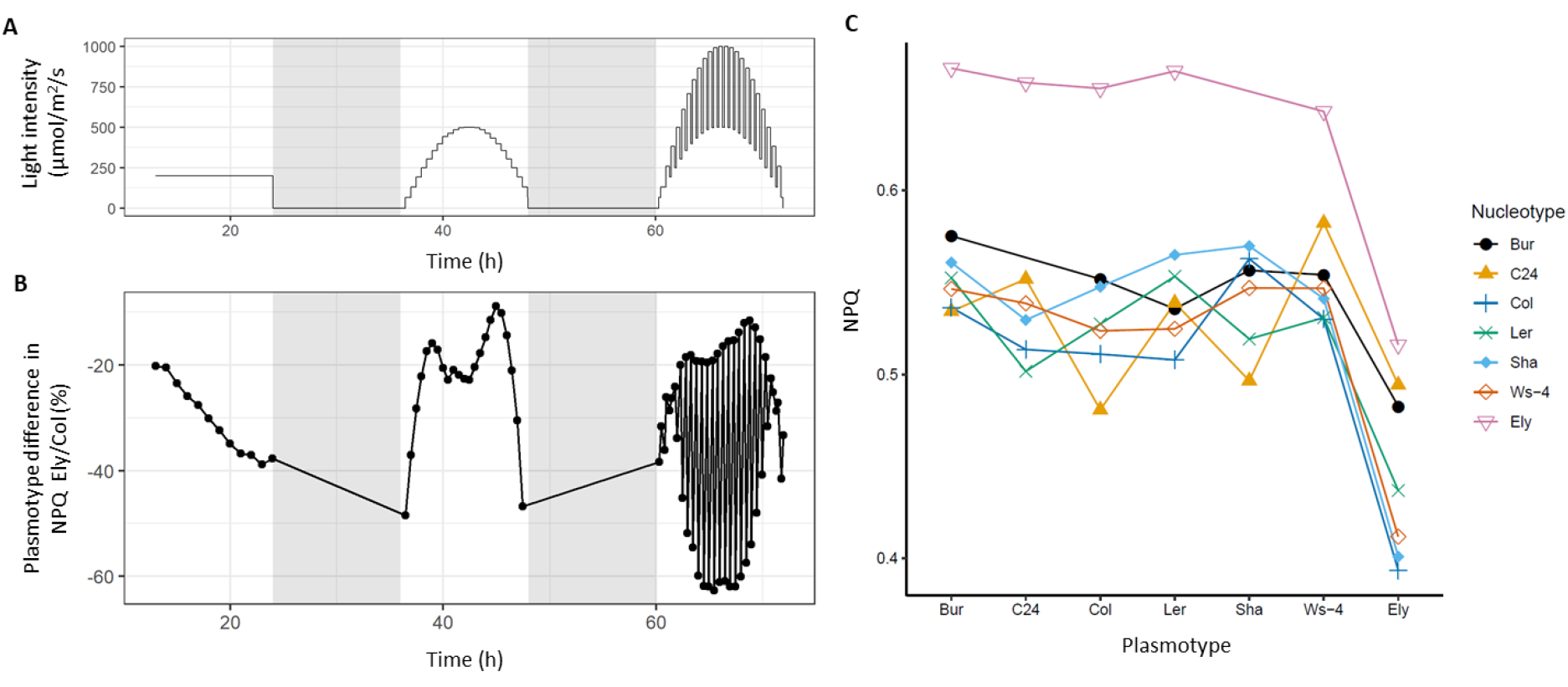
Separating the NPQ phenotypic effects associated with the nucleotype and plasmotype. A) Dynamic light intensity regime as the plants were exposed to, after 21 days growth at 200 µmol m^2^ s^-1^. B) The difference in NPQ between cybrids with the Ely plasmotype compared to the Col plasmotype. The averages were taken over all nucleotypes. All datapoints are negative, meaning the Ely plasmotype has lower NPQ. C) NPQ for all cybrids, at the biggest high to low light transition (at 67 hours in Figure 1A).

To understand the physiological impact of the difference in NPQ between the Ely and Col nucleotype, we further examined the photosynthetic responses under different light conditions. Besides the Ely plasmotype causing the biggest reduction in NPQ in fluctuating light conditions (Figure 1B), also the Ely nucleotype causes significant differences during fluctuating light conditions (Figure 2B and Supplementary Figure 2). During fluctuating light conditions, NPQ is affected most by the Ely nucleotype in comparison to the Col nucleotype in the high to low-light transition compared to the other nucleotypes (Figure 1C). To assess how an increasing light intensity affects the NPQ response after a high to low-light transition, we zoom in to the low light intensity measurements on the first half of the fluctuating light day (from 60 h till 67 h in Figure 2A). Here we observe that the Ely nucleotype has lower NPQ when *Φ*_PSII_ is relatively efficient, whilst the NPQ is higher when *Φ*_PSII_ becomes more inefficient (Figure 2C). Even though this effect is most pronounced after the high to low-light transitions, it is also observable after the low to high-light transitions (Supplementary Figure 3). As NPQ is a measure of capacity, it does not mean that the flux of NPQ (*Φ*_NPQ_) is different. Though we observe that also *Φ*_NPQ_ is higher when *Φ*_PSII_ is less efficient (Figure 2D). This shows that the Ely nucleotype encodes a more efficient NPQ mechanism in fluctuating light conditions.

**Figure 2.**
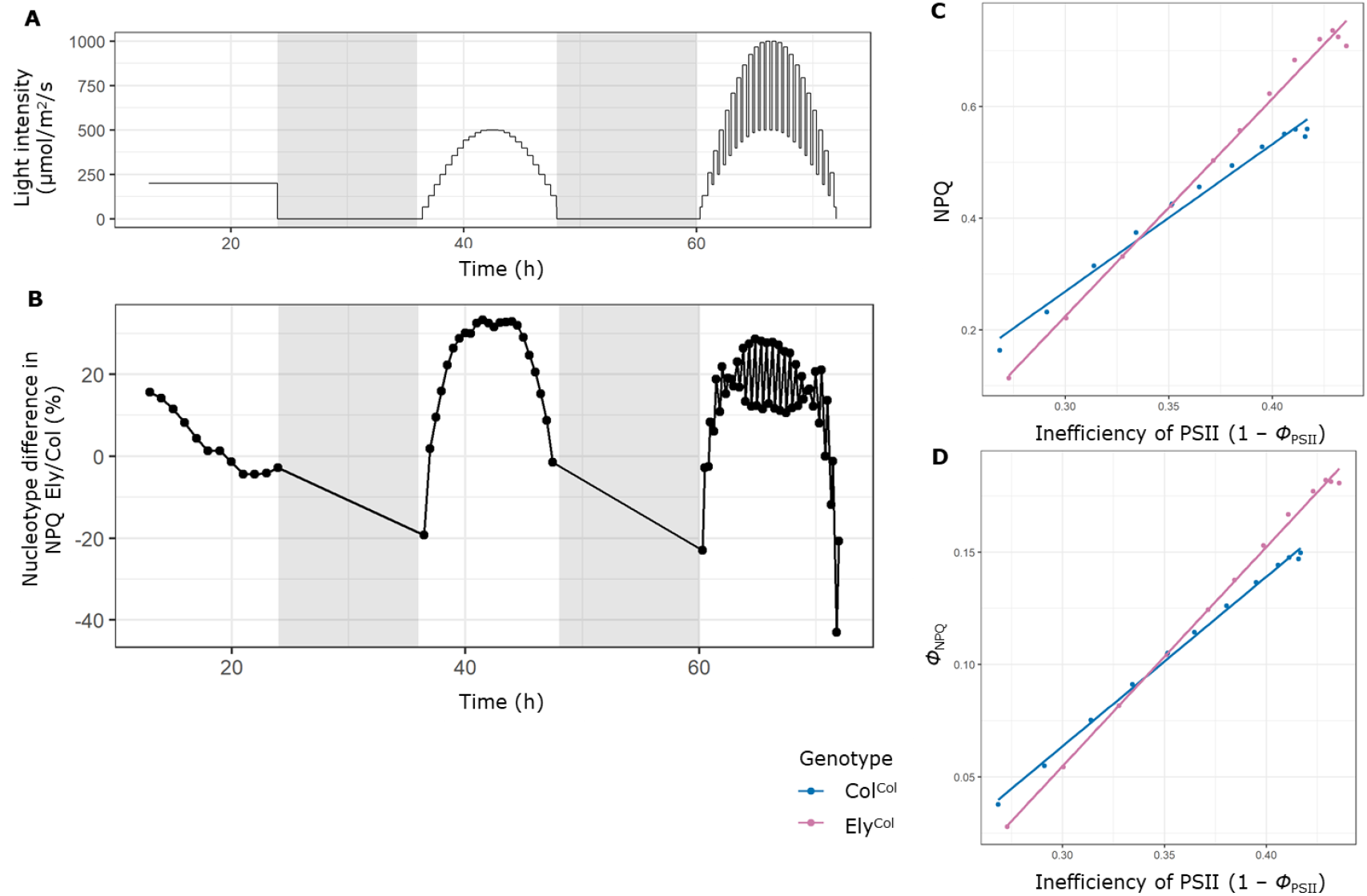
Differences in NPQ capacity and flux between Col and Ely nucleotypes. A) Fluctuating light conditions as the plants were exposed to, after 21 days growth at 200 µmol m^2^ s^-1^. B) The difference in NPQ between cybrids with the Ely nucleotype compared to the Col nucleotype. The average was taken over all plasmotypes. Positive datapoints show the Ely nucleotype has higher NPQ, negative datapoints show the Ely nucleotype has lower NPQ. C) NPQ plotted against the inefficiency of PSII. The measurements are taken 18 minutes after a high to low-light transition, with light intensities as show in panel A from 60 to 67 hours. The low to high-light transition is visualized in Supplementary Figure 3. D) Same as for panel C, but plotted for Φ_NPQ_ plotted against the inefficiency of PSII.

NPQ is determined by the fast-responding component called *q*_E_ and the slow responding component called *q*_I_. To determine which of the two components contributes most to the difference in NPQ, as observed between the Ely and Col nucleotype, we calculate the contributions of each component. The biggest difference was observed after the light transition from 924 µmol m^2^ s^-1^ to 483 µmol m^2^ s^-1^, where the Ely nucleotype shows 28.6% higher NPQ compared to the Col nucleotype. The average NPQ for the Col and Ely nucleotypes at this moment consists of 34.5% and 65.5% of *q*_E_ and *q*_I_ respectively. This shows *q*_I_ is a bigger component of NPQ in these conditions. However, the relative increase in Ely as compared to Col for *q*_E_ is 50.1% while for *q*_I_ this is 18.5%, indicating that *q*_E_ contributes more to the increased NPQ effect induced by the Ely nucleotype. Plotting *q*_E_ and *q*_I_ against the inefficiency of PSII, indeed shows *q*_E_ to resemble the response of NPQ to *Φ*_PSII_ (Supplementary Figure 3). Assessing the ability of the Ely nucleotype to cause faster relaxation of NPQ, via *q*_E_, is difficult with all measurements being taken 18 minutes after a high light intensity period, which is too late to assess the capacity of increased *q*_E_.

To monitor the difference in *q*_E_ between the Ely and Col nucleotypes, we phenotyped the relaxation of NPQ in more detail, after a high to low-light transition. Ely^Col^ and Col^Col^ cybrids were exposed to short fluctuating light conditions (alternating between 1000 µmol m^2^ s^-1^ and 100 µmol m^2^ s^-1^) followed by measuring NPQ during 5 minutes of 50 µmol m^2^ s^-1^. At the end of the high light intensity period (1000 µmol m^2^ s^-1^), Ely^Col^ showed higher NPQ compared to Col^Col^ (Figure 3A). This observation is in line with the earlier observations of higher NPQ due to the Ely nucleotype (Figure 2B). By normalizing the data to the starting NPQ measurement, during the entire 5 minutes NPQ is significantly lower in the Ely compared to Col nucleotypes (Figure 3B). In the first 100 seconds NPQ in Ely shows an average reduction of 13.2% compared to the Col nucleotype (Figure 3B). The experiment was repeated to monitor the NPQ relaxation in a high-light to darkness transition. Ely^Col^ revealed faster relaxation of NPQ in darkness (Figures 3C and 3D), in line with the faster relaxation of NPQ during low light intensities (Figures 3A and 3B). All results together, show that the Ely nucleotype possesses a more efficient NPQ mechanism in conditions where *Φ*_PSII_ is less efficient, and this coincides with the ability to respond more dynamically to fluctuating light conditions as compared to Col.

**Figure 3.**
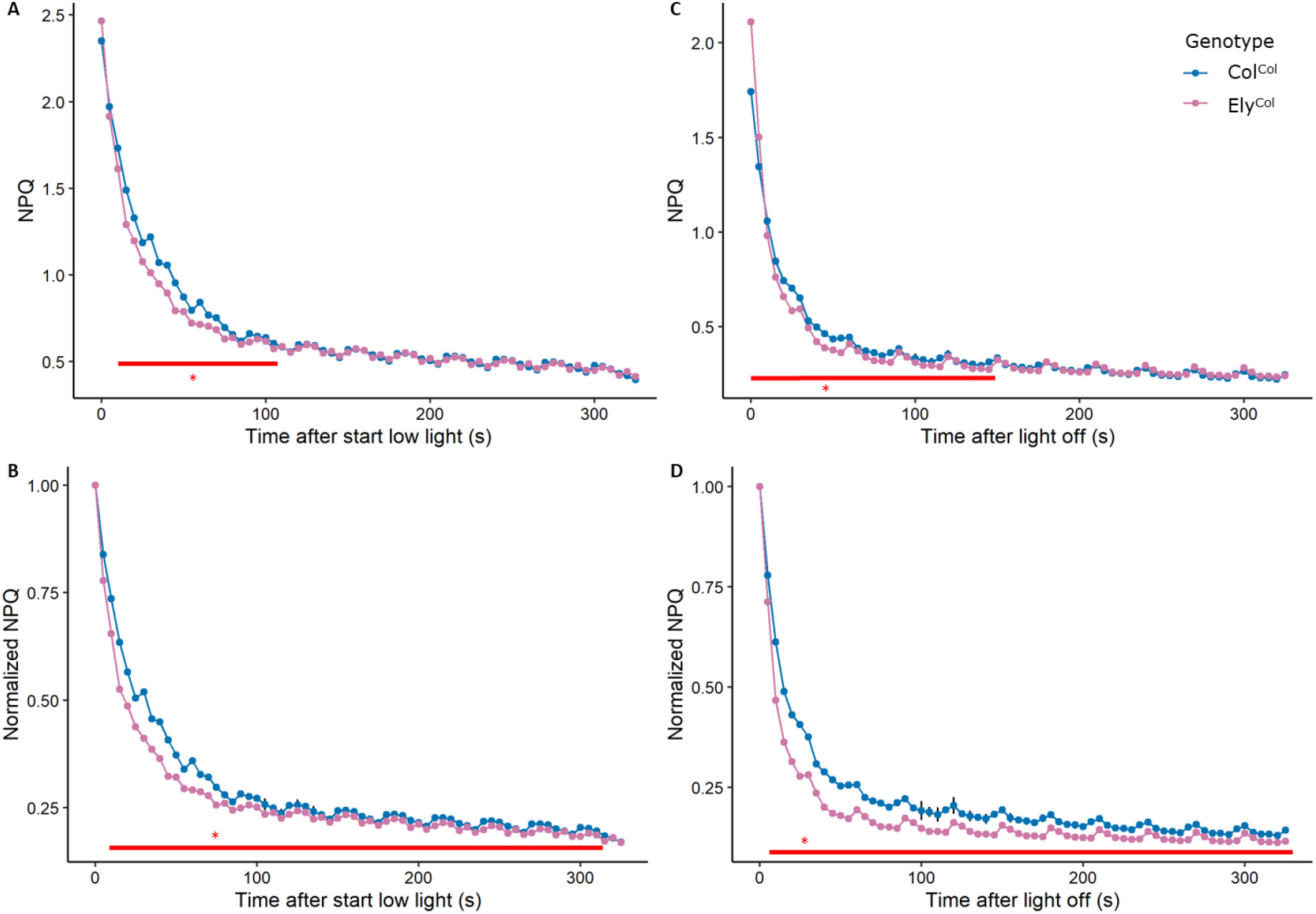
NPQ relaxation of the Ely versus Col nucleotype after fluctuating light conditions. The fluctuating light conditions were five times a three minutes period with 1000, 100, 1000, 100, 1000 µmol m^2^ s^-1^, and at t = 0 s light intensity was switched to 50 µmol m^2^ s^-1^ (shown in panel A and B) and to darkness (shown in panel C and D). Panels A and C show absolute NPQ measurements, while panels B and D show NPQ normalized to the t = 0 s measurement. The red line indicates significant differences between the nucleotypes with n=10.

### Identifying the causal QTLs

Knowing which alleles are causal to the observed difference in capacity and flux of NPQ will help to understand the underlying physiological or biochemical mechanism. NPQ could be higher and more dynamic as a result of the formation of quenching sites in the light harvesting complexes influenced by PsbS and the xanthophyll cycle (Ruban, 2017; Kromdijk and Walter, 2022). The xanthophyll cycle is catalysed by the enzymes VDE and ZEP. We first determined whether there is genetic variation within the genes encoding PsbS, VDE or ZEP, that could explain the observed differences in NPQ. This revealed no non-synonymous variants or impactful INDELs in the genes encoding PsbS, VDE or ZEP. In the absence of genetic variation that may have caused gene expression differences or changes to the protein, we conclude that variant in a different gene (or genes) must be causal.

To identify the causal gene(s) and allele(s) to the difference in NPQ, we constructed a doubled haploid (DH) population between Col^Col^ and Ely^Col^. A DH population is the quickest approach to generate a genetic mapping population of homozygous lines in *A. thaliana*. The population consisting of 449 DH lines segregating between Col^Col^ and Ely^Col^. Low coverage whole genome sequencing and a custom analysis pipeline were used to genotype the DH population (see Materials and Methods). The genotyping resulted in a high resolution marker dataset with 478 markers equally spread over the genome with a resolution of 250 Kbp (Supplementary Data 1). On average six cross overs per DH line were observed (Supplementary Figure 4), so we concluded the DH population formed an excellent starting point to reveal the QTLs involved with the observed difference in NPQ.

To identify the QTLs responsible for the observed differences in NPQ, the DH population was phenotyped using two separate high-throughput chlorophyll fluorescence phenotyping systems. One experiment was designed to provide fluctuating light conditions and the other was designed to provide stable light conditions. In the previous experiment with cybrids, the NPQ difference between the Col and Ely nucleotypes was found to depend on fluctuating light conditions (Figure 2B). The broad sense heritability (H^2^) of the segregating DH population was also found to depend on the light conditions. For NPQ we observed an average H^2^ = 0.25 and in fluctuating light conditions this went up to H^2^ = 0.54, indicating a strong genotype by environment interaction (Supplementary Figure 7). QTL mapping for NPQ throughout the fluctuating light conditions revealed a plethora of QTLs (Figure 4). The majority of QTLs are associated with a specific time of the day, light intensities, sequence of fluctuating light conditions or even adaptation to light conditions. Using a naive Bonferroni threshold (LOD score of 4.8), 15 different QTLs for NPQ can be observed (Figure 4). QTL mapping for *Φ*_PSII_, *Φ*_NPQ_, *Φ*_NO_, *q*_E_ and *q*_I_ identified, next to several shared QTLs, specific QTLs associated with these photosynthetic parameters (Supplementary Figures 8 and 9). In the high-throughput phenotyping system with stable low light conditions (200 µmol m^2^ s^-1^), three QTLs for *Φ*_PSII_ were identified that had not been observed in the fluctuating light experiment (Supplementary Figure 10). Altogether, this shows that there is variation for several physiological mechanisms associated with photosynthetic performance in this population, all of which is dependent on the environmental conditions.

**Figure 4.**
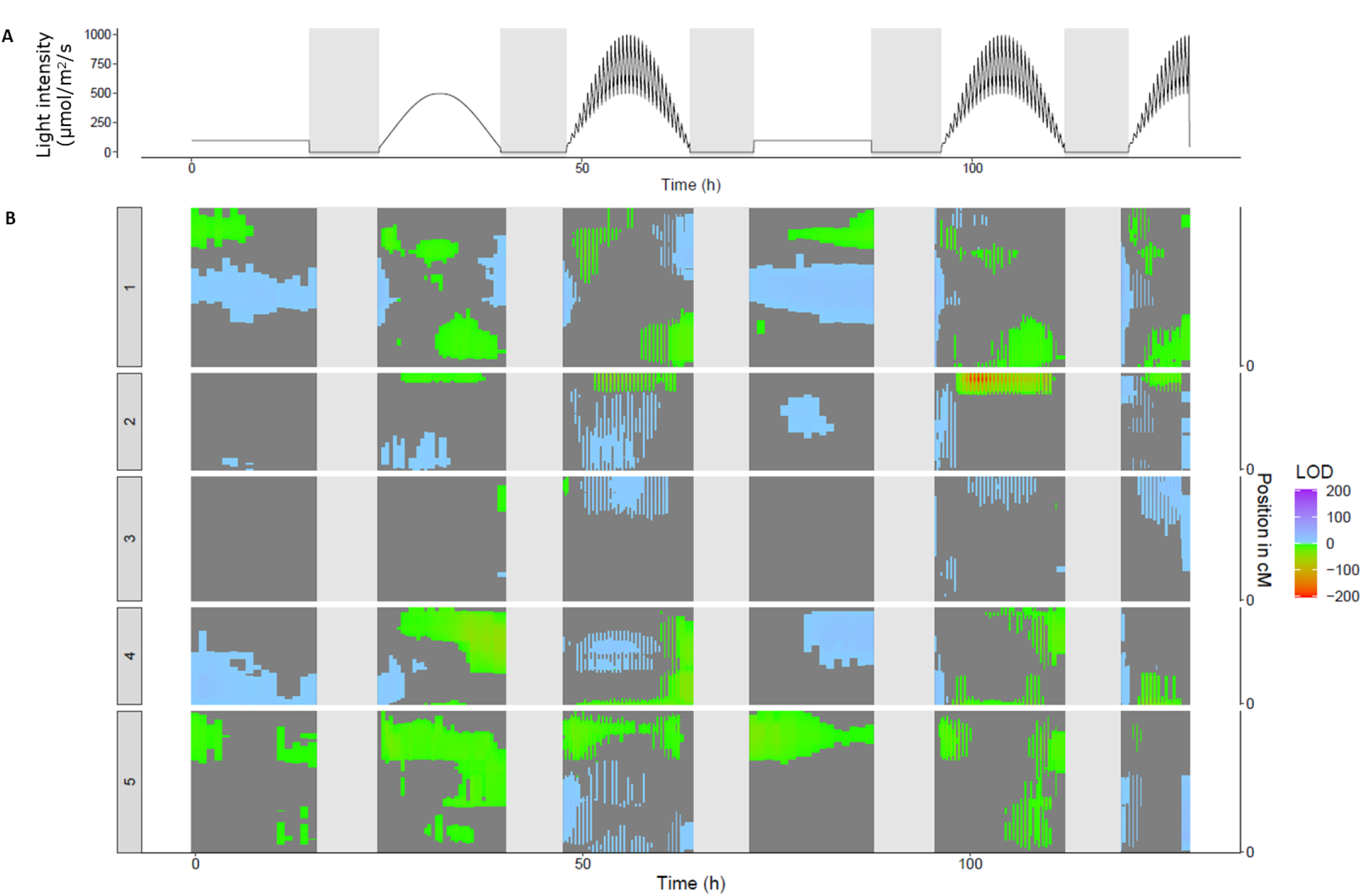
QTL mapping on NPQ using the DH population in both stable and fluctuating light conditions. A) Represents the light intensities during the experiment, where t = 0 h is the moment when lights turned on, day 21 after sowing. In the first 21 days plants were grown at a light intensity of 200 µmol m^2^ s^- 1^. B) Vertical representation of QTL mapping over time, the times match the light intensities are shown in panel A. LOD scores are represented in positive values if the effect size of the Ely allele of a given marker on that time point is higher as compared to Col allele. Negative values are given when the Ely allele induces a lower effect as compared to Col. The dark grey background indicates markers that do not pass a naive Bonferroni threshold (LOD threshold of 4.8).

The results also show that at a specific timepoint for a specific photosynthetic parameter, several QTLs can be observed with opposing effects (Figure 4). As an example, at a single timepoint (t = 86 h) on a day with a stable light intensity (200 µmol m^2^ s^-1^) in between days with fluctuating light conditions, four QTLs for NPQ can be observed (Supplementary Figure 11). Underlying two of these QTLs the Ely allele results in a higher NPQ (a QTL on chromosome 1 at 25.75 Mbp, noted as QTL-1^25,750^, and QTL-5^21,750^ bring about 12.7% and 12.1% higher NPQ respectively). In the other two QTLs the Col allele results in higher NPQ (QTL-1^12,250^ and QTL-4^12,250^ bring about 17.2% and 22.5% higher NPQ respectively). QTLs with opposing effects indicate that the two accessions are likely to use different physiological mechanisms to get to, roughly, the same phenotypic response.

The difference in NPQ as originally observed between the Col and Ely nucleotype was most pronounced in high to low-light transitions (Figure 2). To unravel the underlying genetics, we investigated the two largest contributing QTLs in the fluctuating light conditions. The first QTL is located on chromosome 2 at position 18.5 Mbp, referred to as QTL-2^18,500^. During fluctuating light conditions, QTL-2^18,500^ causes up to 17.5% higher NPQ and 36.3% higher *q*_E_, when homozygous for the Ely allele (Figure 5A). To reveal how the alleles underlying QTL-2^18,500^ affect NPQ at different efficiencies of PSII in fluctuating light, the two parameters were plotted against each other. The Ely allele underlying QTL-2^18,500^ shows an increased capacity of NPQ when PSII is more inefficient (Figure 5D). This resembles the pattern as observed in the parental lines, and we therefore consider QTL-2^18,500^ to explain a substantial part of the differences in NPQ as observed between the Col and Ely nucleotype in the high to low-light transition. Strikingly, QTL-2^18,500^ is absent in low stable light conditions (200 µmol m^2^ s^-1^), meaning that the causal allelic variation does not play a role in such low stable light conditions (Figure 4). During the second day of fluctuating light conditions, the Ely allele of QTL-2^18,500^ has a larger effect, in comparison to the first fluctuating light day, implying that the underlying mechanism is adapting to the environmental conditions (Figure 6B). In the high to low-light transition when the NPQ effect is largest, *Φ*_PSII_ is reduced with 3.4%, which is in line with higher NPQ when PSII is less efficient (Figure 6C).

**Figure 5.**
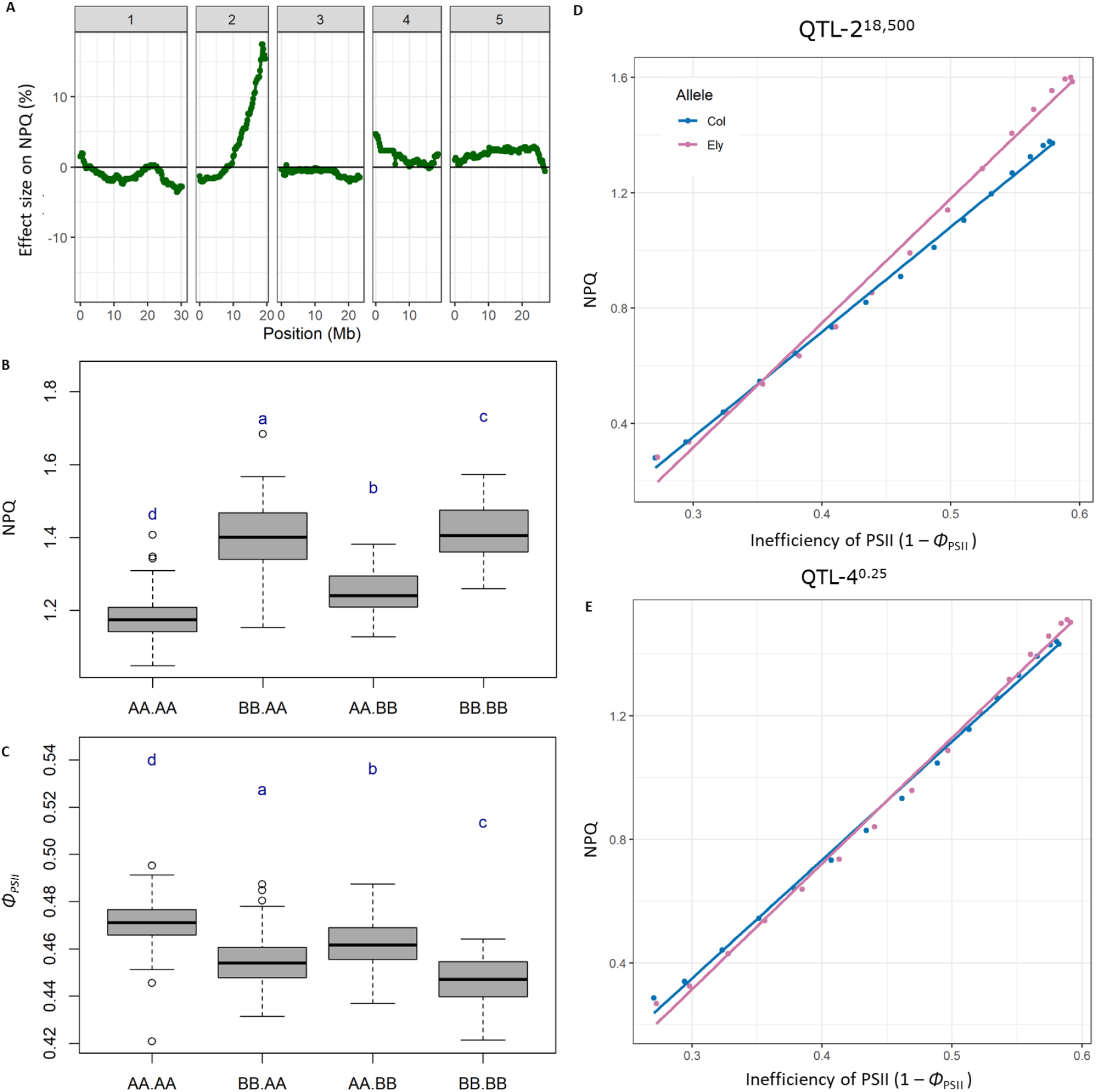
Phenotypic effects of QTL-2^18,500^ and QTL-4^0.25^ on Φ_PSII_ and NPQ. A) Shows what the effect of the Ely or Col allele at a certain position on the A. thaliana genome is on NPQ. The effect size is normalized to the Col allele, so a positive effect size is higher NPQ conferred by the Ely allele. The x axis shows the position (in Mbp) on the five chromosomes of A. thaliana. B and C) Phenotypic effects of the alleles underlying QTL-2^18,500^ and QTL-4^0.25^ on Φ_PSII_ and NPQ. The phenotypes are given at the timepoint when the difference between the Ely and Col nucleotype was largest, in the middle of the second fluctuating light day, at 101 h into the experiment (Figure 4A). The phenotypic effects are given for all homozygous combinations between QTL-2^18,500^ and QTL-4^0.25^, with the two left letters representing QTL-2^18,500^ and the two right letters representing QTL-4^0.25^. The “A” allele refers to the Col allele, and the “B” allele refers to the Ely allele of both QTLs. The letters represent significant differences, with Tukey correction for multiple testing (α = 0.05). D) The NPQ effect of the Col and Ely allele underlying QTL-2^18,500^ plotted against the inefficiency of PSII. The measurements are taken 18 minutes after a high to low-light transition. The measurements are taken from the second day with fluctuating light conditions, from 96 to 102 h (Figure 4A). E) Same as panel D, but for the alleles underlying QTL-4^0.25^.

**Figure 6.**
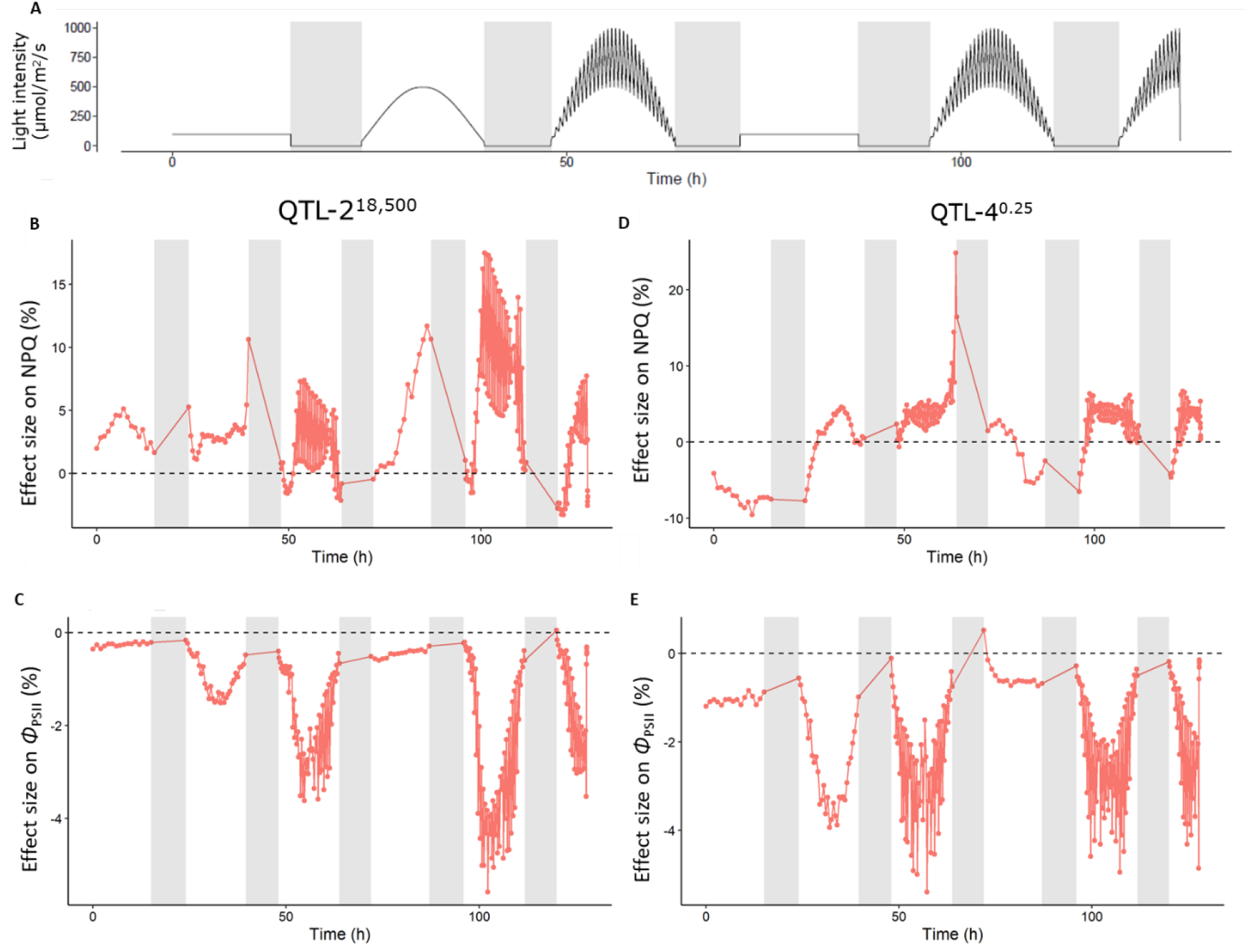
Phenotypic effect sizes of QTL-2^18,500^ and QTL-4^0.25^ on Φ_PSII_ and NPQ. A) Represents the light intensities during the experiment, where t = 0 h is the moment when lights turned on, day 21 after sowing. In the first 21 days plants were grown at a light intensity of 200 µmol m^2^ s^-1^. B) Effect size of alleles underlying QTL-2^18,500^ on NPQ with Col as reference. C) Effect size of alleles underlying QTL-4^0.25^ on NPQ with Col as reference. D) Effect size of alleles underlying QTL-2^18,500^ on Φ_PSII_ with Col as reference. E) Effect size of alleles underlying QTL-4^0.25^ on Φ_PSII_ with Col as reference. In panels A – E) the effect size matches the light intensities shown in panel A.

At the time point at which the difference between the alleles underlying QTL-2^18,500^ is the largest, the Ely allele of another QTL confers 4.7% higher NPQ than the Col allele. This QTL is located on chromosome 4 at position 250 Kbp, and is referred to as QTL-4^0.25^ (Figure 5A). QTL-4^0.25^ is present in stable low light conditions as well as fluctuating light conditions (Figure 4 and Supplementary Figure 10). Throughout the experiment, the Ely allele can be seen to confer a difference in NPQ ranging between 24.8% higher and 9.5% lower compared to the Col allele (Figure 6D). Also, *Φ*_PSII_ is found to depend on the light conditions, with the Ely allele conferring up to 5.4% lower *Φ*_PSII_ compared to the Col allele (Figure 6E). However, both alleles underlying QTL-4^0.25^ at any given *Φ*_PSII_ have the same NPQ phenotype (Figure 5E), in contrast to QTL-2^18,500^, that did show a difference in NPQ (Figure 5D). Analysing the combined effect of the alleles underlying the two QTLs, shows that for *Φ*_PSII_, both QTLs have an equal effect, regardless of the alleles underlying the other QTL (Figure 5C). Doing this for NPQ; when a plant has the Ely allele underlying QTL-2^18,500^, the presence of the Ely allele underlying QTL-4^0.25^ increases NPQ only by 0.6%. While doing the same conversion with the Col allele for QTL-2^18,500^, alleles underlying QTL-4^0.25^ caused a 6.2% increase (Figure 5B). Therefore, for NPQ the alleles underlying the two QTLs are in an epistatic relation (p = 8.21E-5). This shows that the physiological processes caused by the alleles underlying the two QTLs are independent.

### Fine mapping and gene confirmation

To reveal the physiological mechanisms conferred by the alleles underlying the two QTLs, the causal genes should be identified. Thus, we performed various fine mapping approaches and gene candidate validations for both QTLs. QTL-2^18,500^ showed the highest association at 18.5 Mbp on chromosome 2, but due to the high association and relatively large linkage disequilibrium, the QTL is wide (Figure 7A). On both sides of QTL-2^18,500^, we used the recombination sites within two individual DH lines that were closest to the marker, to determine the size of the QTL. Using this definition, QTL-2^18,500^ stretched from 18,612,500 to 18,862,500 bp – i.e the size is 250 Kbp. To remove other QTLs that can interfere with the phenotype of this QTL when doing fine mapping, we produced a range of 57 near-isogenic lines (NILs). All NILs had independent recombinations in the QTL region of 250 Kbp. These recombinants were phenotyped in the same way as the original DH population, and the QTL mapping showed the highest association on 18,861,385 bp (Figure 7B). Again, using the recombination sites within two individual fine mapping lines, on both sides, closest to 18,861,385 bp, showed the region to span from 18,849,338 bp to 18,875,047 bp which is 25 Kbp in size.

**Figure 7.**
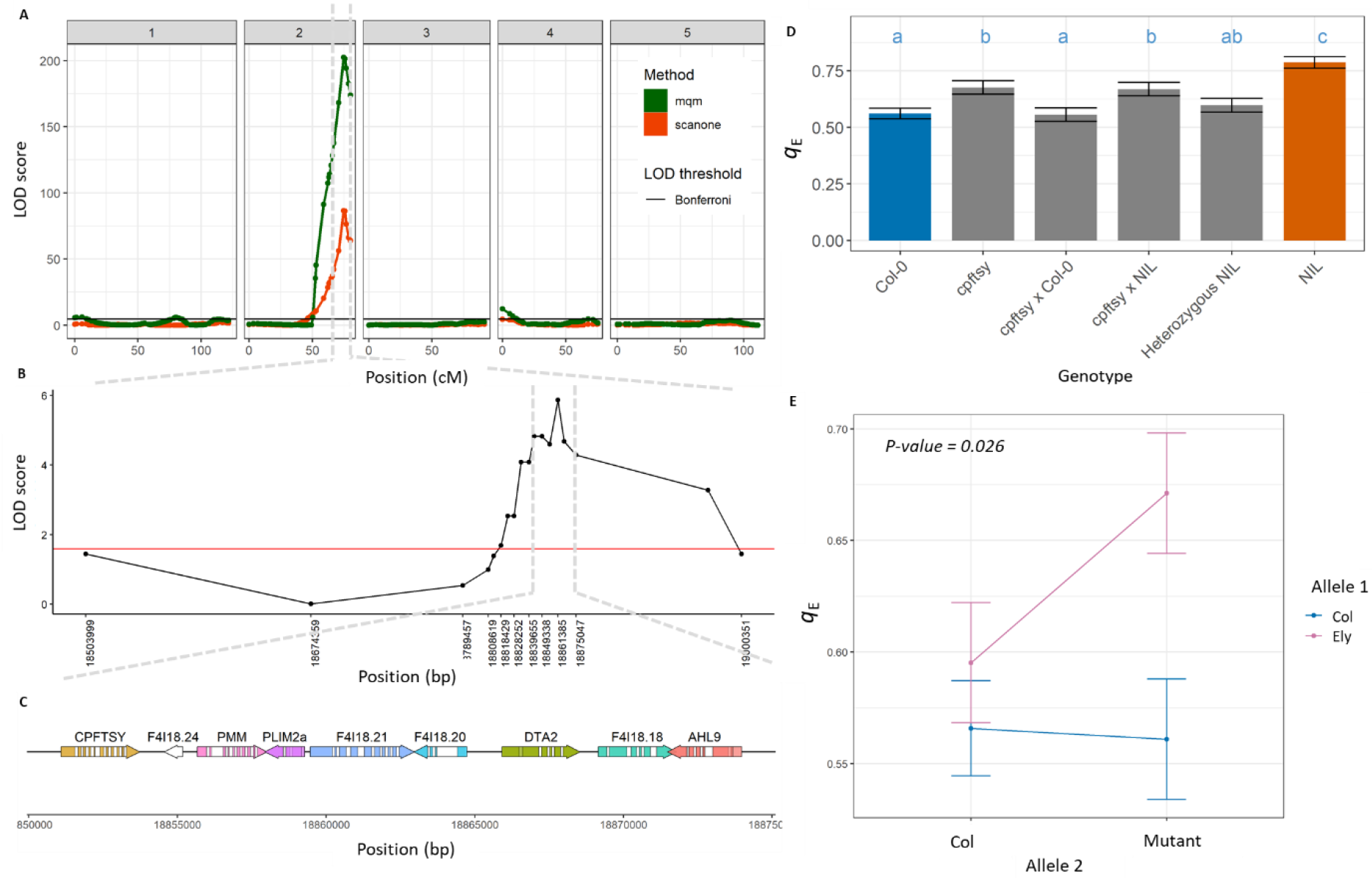
QTL mapping, fine mapping and quantitative allelic complementation studies to identify the causal genes underlying QTL-2^18,500^. A) The QTL map of the DH population for NPQ during fluctuating light at timepoint 101 h after the start of the experiment. The LOD threshold is based on a naive Bonferroni correction, here established at 4.8. B) The QTL map of the fine mapping population, consisting of 57 recombinants in a 250 Kbp region around the QTL. C) Genes annotated within the QTL region of 25 Kbp. D) q_E_ 102 h into the experiment, of Col-0, cpftsy and the NIL, and F_1_ hybrids between these. The NIL is coloured orange in line with Figure 9. E) Quantitative allelic complementation identifying cpFtsY is the causal gene underlying QTL-2^18,500^. Allele 1 is either the Col or knock-out allele, and Allele 2 is either the Col or Ely allele, in the F_1_ hybrids. In all cases significance is displayed as letters, with a Tukey multiple correction (α = 0.05).

There are nine genes annotated within the 25 Kbp region that contains the causal gene (Figure 7C). We analysed the genetic variation present within this region to find if any variant is expected to cause a difference in protein abundance or structure. Variant prediction and *de novo* sequencing of Ely shows four non-synonymous SNPs, one frameshift and two promotor deletions in six of these genes (Supplementary Table 1). A non-synonymous SNPs is a SNP in the predicted open reading frame of the gene, and a frameshift is an insertion or deletion that influences the reading frame, and both are expected to lead to a change in amino acid(s) of the encoded protein. To assess the effect of the promotor deletions on expression of the genes, we compared the expression of *PHOSPHOMANNOMUTASE* (*PMM*) and *CHLOROPLAST SIGNAL RECOGNITION PARTICLE FTSY* (*cpFtsY*) between Col and Ely, before and during fluctuating light (Supplementary Figure 14). This revealed no significant expression differences between Col and Ely for *PMM*. In the absence of a non-synonymous SNP this makes *PMM* an unlikely causal gene. Also, *cpFtsY* did not show a significant expression difference between Col and Ely. However, the presence of a non-synonymous SNP, and the previously determined role in incorporating light harvesting complexes into the thylakoid membrane (Durrett *et al*., 2006; Tzvetkova-Chevolleau *et al*., 2007), lead us to test this gene for causality first. A T-DNA insertion knock-out line of *cpFtsY* showed increased *q*_E_, although not as high as the NIL for this QTL showed (Figure 7D). The genotype that is heterozygous for the remaining 25 Kbp region shows the same *q*_E_ phenotype as the Col wildtype control, meaning the Ely allele is recessive (Figure 7D). Due to the Ely allele being recessive, we can test the complementary effect of the two different alleles in an F_1_ hybrid with the T-DNA insertion knock-out line, a test called quantitative allelic complementation. In these F_1_ hybrids, we saw that the Ely allele is not able to complement the *cpFtsY* knock-out mutant phenotype, as seen with the Col allele, hence we can conclude that *cpFtsY* is the likely causal gene explaining the increased NPQ capacity when *Φ*_PSII_ is inefficient (Figure 7E).

For QTL-4^0.25^, we had to take into account the, well-studied, inversion on top of chromosome 4 (Fransz *et al*., 2016) (Figure 8A). Using the genetic map and the *de novo* sequence of Ely we could establish that QTL-4^0.25^ is 1.25 Mbp away from this inversion (Supplementary Figures 5 and 12), and therefore can be assessed without taking the inversion into account. To determine the size of QTL-4^0.25^, we used the recombination sites within two individual DH lines that were closest on both sides of the highest associated marker. This resulted in a QTL spanning from 237.5 to 337.5 Kbp, meaning the causal gene is located within a 100 Kbp region. Three NILs were made with different recombination sites within the 100 Kbp region (Figure 8B). Phenotyping these NILs for *Φ*_PSII_ and using the recombination sites, showed that the QTL is located between 260,617 bp to 324,150 bp, making the region containing the QTL 64 Kbp in size (Figure 8B).

**Figure 8.**
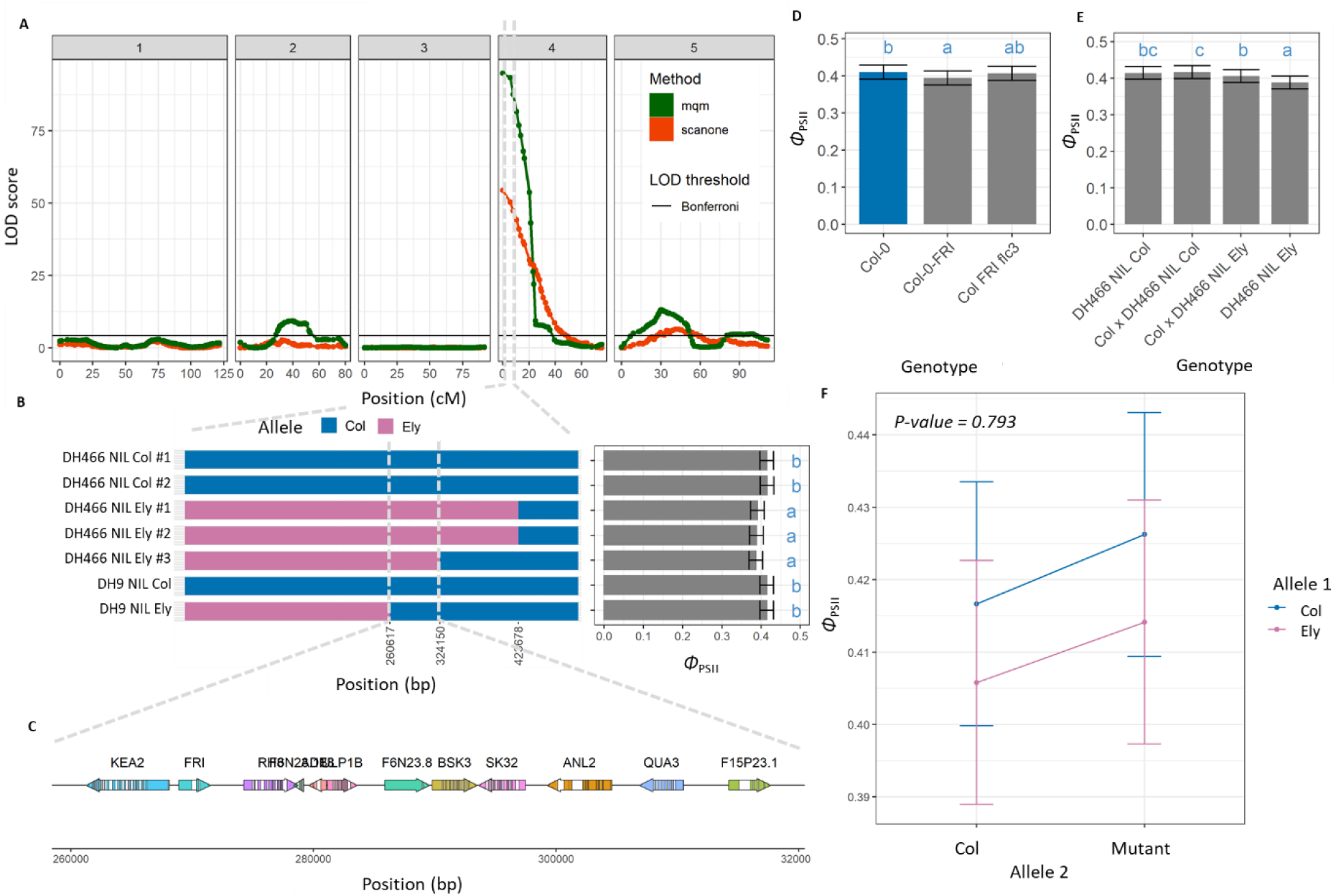
QTL mapping, fine mapping and quantitative allelic complementation studies to identify the causal genes underlying QTL-4^0.25^. A) The QTL map of the DH population for Φ_PSII_, 1 h into a high light intensity of 500 µmol m^2^ s^-1^, after plants were grown for 18 days at 200 µmol m^2^ s^-1^. LOD threshold is based on a naive Bonferroni correction, here placed at 4.1. B) The genotype of separate near-isogenic lines and their controls within the QTL region, and the associated Φ_PSII_ phenotypes. C) Genes annotated within the QTL region of 64 Kbp. D) Phenotype of Col genotypes with different alleles for FRI and FLC, at the same time point as the original QTL mapping. E) Phenotypes of heterozygous NILs in comparison to the controls. F) Quantitative allelic complementation study for the KEA2 gene. Allele 1 is either the Col or knock-out allele, and Allele 2 is either the Col or Ely allele, in the F_1_ hybrids. In all cases significance is displayed as letters, with a Tukey multiple correction (α = 0.05).

There are 11 genes annotated within the 64 Kbp region surrounding QTL-4^0.25^ (Figure 8C). With the highest association in the QTL mapping of the DH population being at 267,500 bp, we first focused on the genes around this position. Position 267,500 bp is located 1,401 bp upstream of the annotated position of the flowering time gene *FRIGIDA* (*FRI*). Ely has an active allele and Col has a knock-out allele of *FRI*, meaning these alleles segregate within the DH population. *FRI* is known for many pleiotropic effects, and thus the allelic differences between Col and Ely could explain the difference in *Φ*_PSII_. To test the role of *FRI*, we used a late-flowering Col genotype with the functional allele of *FRI* introgressed from Sf-2 (i.e. Col-*FRI*) (Lee *et al*., 1993), and Col genotype with an active allele of *FRI* that is early flowering due to an *flc* knock-out (i.e. Col-FRI-flc3). Noteworthy, a biparental population between Col and Sf-2 did not show a QTL on chromosome 4 for NPQ, already suggesting that the difference between a knock-out and functional allele of *FRI* are not causing the observed phenotypic difference (Jung and Niyogi, 2009). Phenotyping for *Φ*_PSII_ showed that the Col-*FRI* genotype showed a small significant effect in comparison to the Col genotype (Figure 8D). However, in the absence of significant effect between the Col-*FRI-flc3* and the Col genotype, shows that the active allele of *FRI* cannot explain the phenotypic difference as caused by the alleles underlying QTL-4^0.25^ (Figure 8D). Independently of this, we phenotyped three recombinant inbred line (RIL) populations derived from the crosses between Can-0 and Col, Sha and Col and Bur-0 and Col. Bur-0, Sha and Can-0 all harbour the functional allele of *FRI* (Shindo *et al*., 2005; Brachi *et al*., 2010), and thus if a functional *FRI* allele would explain the difference in *Φ*_PSII_, a QTL would be present on the top of chromosome 4 for all three populations. As the RILs with Bur and Sha do not show a QTL on chromosome 4 (Supplementary Figure 13), we can exclude *FRI* as the causal gene of QTL-4^0.25^. Based on a gene function analysis, the other likely candidate gene would be the *K+ efflux antiporter 2* (*KEA2*). However, the quantitative allelic complementation using the NILs showed that both alleles of QTL-4^0.25^ were able to complement the mutant in the same way (Figure 8F). This does not exclude *KEA2* as the causal gene, but further experiments would have to prove it.

### Physiological impact of the alleles

To understand how the observed differences in NPQ are caused by the studied QTLs, we sought to connect the gene function of the causal genes with more in-depth physiological insights. As for now we only know the causal gene for QTL-2^18,500^, we primarily focus on the physiological impact of the gene *cpFtsY*. The earlier observations that the Ely allele underlying QTL-2^18,500^ results in higher engagement of NPQ when *Φ*_PSII_ is less efficient, can now be connected to the Ely allele of *cpFtsY* (Figures 2C and 2D). Assessing the effect of *cpFtsY* at different NPQ intensities shows that the Ely allele has lower 1 – q_P_ when NPQ is higher (Figures 9A and 9B). A lower 1 – q_P_ means the Q_A_ pool is more oxidized, which could lead to less damage to the reaction centra, and also less chlorophyll triplets being formed in the pigment bed.

**Figure 9.**
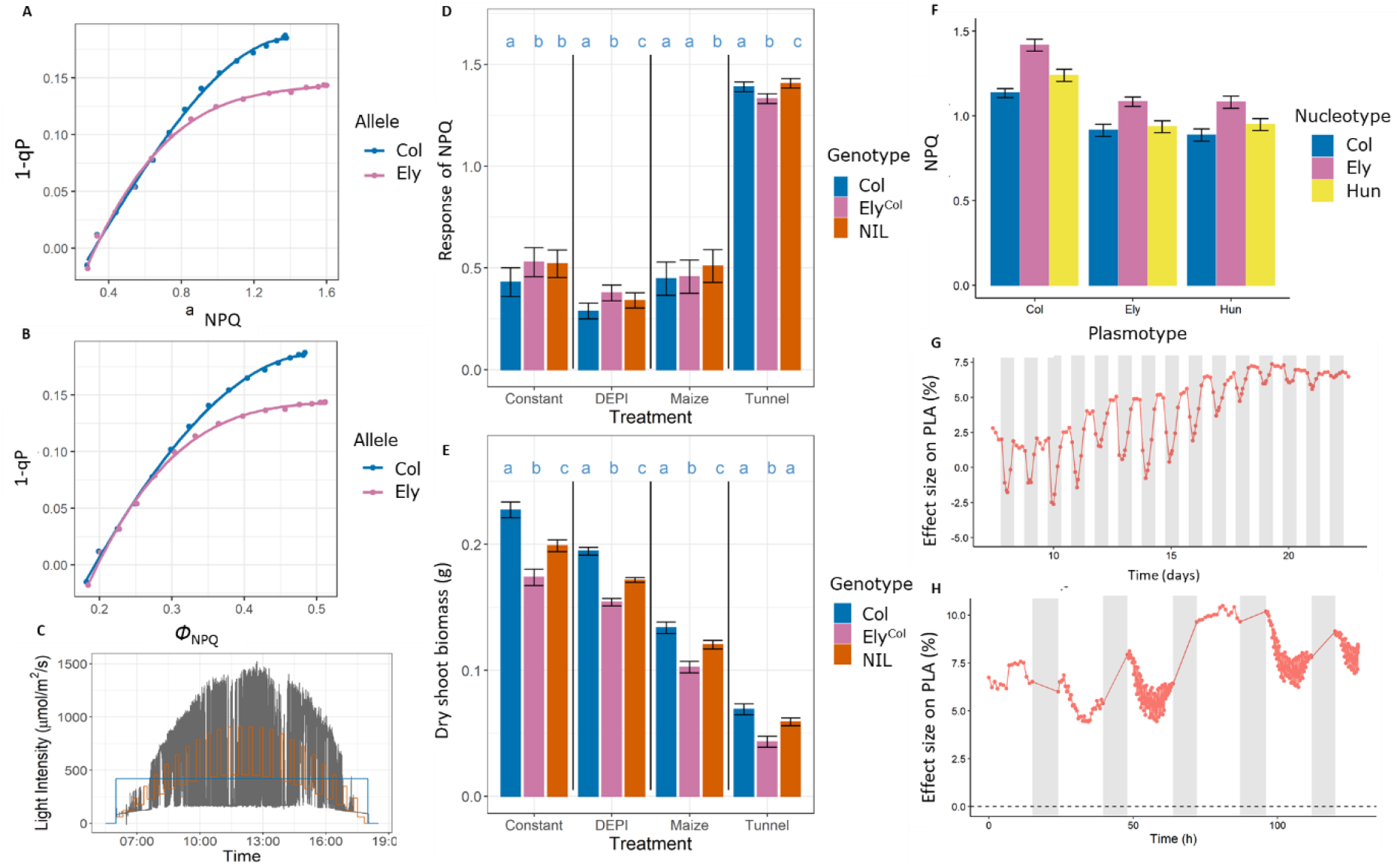
Assessment of the physiological impact of the Col and Ely alleles of QTL-2^18,500^ and QTL-4^0.25^. A) Capacity of NPQ (NPQ) plotted against Q_A_ (1 - q_P_) during the low light periods in fluctuating light conditions for QTL-2^18,500^. The measurements are taken 18 minutes after a high to low-light transition. The measurements are taken from the second day with fluctuating light conditions, from 96 to 102 h (Figure 4A). B) The same as panel A, but for the flux of NPQ (φ_NPQ_) against Q_A_ (1 - q_P_). C) The three light intensity treatments the genotypes were exposed to on a daily base. In blue the stable light condition is shown (named “Constant” in panels D and E), in orange the sinusoidal fluctuating light condition is shown (named “DEPI”) and in grey the highly fluctuating light condition is shown (named “Maize”). In all three conditions the photoperiod and average light intensity is the same. The light intensity for the plants grown in the semi-protected tunnel is shown in Supplementary Figure 15 (named “Tunnel” in panels D and E). D) The response of NPQ for the Col^Col^, Ely^Col^ and NILs possessing QTL-2^18,500^ grown in different light regimes, as a ratio between NPQ before and after a 2 minute high light pulse of 1000 µmol m^2^ s^-1^. Significance is displayed as letters, within one treatment, with a Tukey multiple correction (α = 0.05). E) The dry shoot biomass (g) for the genotypes as exposed to the four different light treatments 28 days after sowing. Significance is displayed as letters, within one treatment, with a Tukey multiple correction (α = 0.05). F) Cybrids with all combinations between the nucleotypes and plasmotypes between Col-0, Ely and Hun-0 measured in the same way as the fluctuating high-throughput phenotyping experiments as used for the original cybrids, DH mapping and fine mapping experiments. NPQ given at the time point where the NPQ difference between the Ely and Col nucleotype was largest. G) Plant leaf area measurement for the QTL-4^0.25^ in the DH population as effect size of the Ely allele in comparison to the Col allele, where a positive value indicates larger plant leaf area caused by the Ely allele. The plant leaf area was tracked for the full 23 days of the stable light experiment. H) Same as in panel G, but during the fluctuating light experiment, where t=0 depicts the morning of day 21 of after sowing. The full 21 days the conditions were identical to plants grown in the stable light experiment as shown in panel G.

Assessing the impact of higher engagement of NPQ on overall plant performance was done by growing three independent NILs containing the Ely allele underlying QTL-2^18,500^ in the Col background, as well as Col^Col^ and Ely^Col^, in four different environmental conditions. As the Ely *cpFtsY* allele showed a higher NPQ response especially in high and fluctuating light conditions, but not during low light conditions, we grew these lines in stable low light conditions, sinusoidal fluctuating light conditions, highly fluctuating light conditions and in a semi-protected tunnel in the spring of 2021 (Figure 9C). We observed that indeed NPQ (measured as NPQ_t_) was higher in the NILs compared to the Col^Col^ control (Figure 9D). This measure of NPQ is expressed as the ratio of NPQ after a short high light exposure (two minute high light pulse of 1000 µmol m^2^ s^-1^), in comparison to the low light conditions before high light intensity. The higher NPQ effect is observed in all treatments, and thus seems to be independent of the growing conditions (Figure 9D). Likewise, the above ground biomass of the NILs is less than in the Col control (Figure 9E). Thus, the Ely *cpFtsY* allele results in lower biomass, in all tested conditions, in comparison to the Col allele.

A Ser-264-Gly amino acid substitution in the Ely *PsbA* allele does not only confer resistance to triazine herbicides, it also reduces the affinity of the Q_B_ site for plastoquinone (Oettmeier, 1999). We already observed that the Ely *PsbA* allele results in reduced *Φ*_PSII_ (Figure 1B). Therefore, higher engagement of NPQ when *Φ*_PSII_ is less efficient may be considered as a compensation mechanism in the Ely wild type accession, with the *PsbA* mutation. Whether this is a requirement for the Ely accession to survive in its natural habitat is difficult to assess. (Flood *et al*., 2016*a*) found the *PsbA* mutation to be wide spread along the British railway system. However, the nucleotypes of all accessions found to have the *PsbA* mutation were nearly identical to Ely, making it difficult to draw conclusions on. Meanwhile, we managed to find one accession, originating from Huntly railway station (Huntly, Scotland, UK; Hun-0), which carries the same *PsbA* mutation as found in Ely, but in an unrelated nucleotype background. Like Ely, this is a largely homozygous accession, likely resulting from multi-generation inbreeding. This makes it very likely that Hun-0 carries a *PsbA* mutation that arose independently from the *PsbA* mutation first identified in Ely. Comparing the Hun^Col^ cybrid with the Col^Col^ and Ely^Col^ cybrids, shows that the Hun-0 nucleotype does not confer the same compensatory mechanism as conferred by the Ely nucleotype (Figure 9F). This suggests that for plants to survive on the railways, the Ely nucleotype compensatory mechanism is not required when a *PsbA* mutation is present.

The causal gene underlying QTL-4^0.25^ for now remains unknown. Nevertheless, the Ely allele for QTL-4^0.25^ contributes to a 7.5% larger projected leaf area, as measured repeatedly in different experimental set-ups (Figures 9G and 9H). This positive effect on plant leaf area is either contributed to the causal gene conferring the difference in NPQ and *Φ*_PSII_, or to a gene very closely linked to it. This as the late flowering induced by the *FRI* locus is known for pleiotropic effects, and larger plant leaf area could well be one of such pleiotropic effects. Assessing the link between the causal gene and biomass remains to be done, once the causal gene is identified.

## Discussion

### Revealing a nucleotype with faster relaxing NPQ

The *A. thaliana* accession Ely is known to have a large effect mutation in the plasmotypic *PsbA* gene (El-Lithy *et al*., 2005; Flood & Theeuwen *et al*., 2020). Here, we discovered that also the nucleotype of Ely conveys large photosynthetic differences, compared to six other *A. thaliana* accessions. This finding was possible due to the ability of distinguishing phenotypic differences caused by the plasmotype, from differences caused by nucleotype (Flood & Theeuwen *et al*., 2020). During fluctuating light conditions, the Ely wildtype accession had a roughly equal NPQ phenotype compared to the Col wildtype accession. With the Ely plasmotype reducing NPQ significantly, this showed that the Ely nucleotype was able to increase NPQ. Compared to the Col nucleotype, the Ely nucleotype causes lower NPQ when PSII is highly efficient and higher NPQ when PSII is highly inefficient. (Figure 2). The difference in NPQ between the nucleotypes was primarily caused by the fast relaxing component, *q*_E_. Faster relaxing NPQ has been defined as a desirable photosynthetic property, as it is one of the bottlenecks limiting photosynthesis in crops (Zhu *et al*., 2004, 2010). NPQ relaxation experiments showed that the Ely nucleotype has ability to relax NPQ faster than the Col nucleotype (Figure 3). Compared to the Col nucleotype, the Ely nucleotype has higher NPQ at the switch from high light to low light intensity. However, the Ely nucleotype reduced NPQ faster than the Col nucleotype, this was apparent within 10 seconds of the light switch and the NPQ remained lower for up to 150 seconds. When normalized against the starting point, NPQ was found to be significantly lower during the entire period of low light intensity, showing the relative ability of the Ely nucleotype to relax NPQ faster. Tobacco and soybean that were genetically modified to have faster NPQ relaxation showed a similar pattern of relaxation (Kromdijk *et al*., 2016; De Souza *et al*., 2022). In these crops, this faster NPQ relaxation was shown to result in higher yields.

### Plethora of QTLs influencing NPQ in dynamic light conditions

Faster NPQ relaxation in tobacco and soybean was achieved by overexpressing the genes encoding PsbS, VDE and ZEP (Kromdijk *et al*., 2016; De Souza *et al*., 2022). Earlier QTL mapping approaches have revealed genetic variation in *PsbS* to be causal to phenotypic variation in NPQ (as found by Wang *et al*., 2017*b*; Rungrat *et al*., 2019). However, no genetic differences within the promotor regions or genes encoding PsbS, VDE and ZEP were found between the Ely and Col nucleotypes. To reveal the underlying genetic architecture, a doubled haploid population between Ely and Col was constructed. Phenotyping the segregating population in dynamically controlled high-throughput phenotyping systems revealed variable broad sense heritability, depending on the light conditions. Variability in heritability is commonly caused by gene by environment interactions (Visscher *et al*., 2008). As NPQ is a dynamic process and commonly upregulated in high light and fluctuation light conditions (Long *et al*., 2022), genetic variation resulting in differences in NPQ may be most pronounced in such environmental conditions. Mapping the response of NPQ in the DH population to fluctuating light conditions revealed not just one QTL responsible, but a plethora of QTLs (Figure 4).

The QTL mapping of other photosynthetic traits, *Φ*_PSII_, *Φ*_NPQ_, *Φ*_NO_, *q*_E_ and *q*_I_ identified additional QTLs. Many of the identified QTLs showed to be highly dependent on specific light conditions. Time of day, adaptation and duration of a light period all influences whether a QTL affects NPQ. The results also show that several QTLs can be observed with opposing effects at specific time points for a specific photosynthetic parameter (Figure 4). Some of the QTLs showed opposing effect depending on the allele. This indicates that both accessions use different physiological mechanisms to get to roughly the same phenotype. While normally implicitly assumed that differences in NPQ are primarily caused by expression of PsbS, VDE and ZEP, the absence of these genes underlying the QTLs detected here, shows that allelic variation in many more genes can result in photosynthetic variation. Oakley *et al*., 2018 showed how phenotyping a different biparental mapping population in *A. thaliana* in cold stressed plants also revealed a range of different QTLs for NPQ. Therefore, we conclude that genotypes can contain a range of allelic variants that result in different photosynthetic responses to different and changing environmental conditions, as compared to other genotypes.

In this study a biparental mapping population was chosen to identify the causal genetic variation underlying differences in NPQ between the two accessions. While depending on the parental lines, the many QTLs identified here and by Oakley *et al*., 2018 show that biparental mapping populations have strong statistical power to reveal many QTLs. To determine whether all QTLs for NPQ present within the population were identified would require calculating percentage variance explained by the detected QTLs (Kruijer *et al*., 2015), but because of the many QTLs involved, more accurate methods for these calculations need to be explored (Kruijer, personal communication). GWAS are often used as an alternative to biparental mapping populations to reveal genetic variation for photosynthesis (Bezouw *et al*., 2019). GWAS have as advantage that the population contains more genetic diversity and lower linkage disequilibrium (Korte and Farlow, 2013). However, GWAS for photosynthetic phenotypes generally identified less QTLs (Chao *et al*., 2014; Ortiz *et al*., 2017; Wang *et al*., 2017; Van Rooijen *et al*., 2017; Rungrat *et al*., 2019; Prinzenberg *et al*., 2020; Joynson *et al*., 2021; Ferguson *et al*., 2021). This is likely the result of lower statistical power caused by rare alleles with small effect sizes (Korte and Farlow, 2013; Forsberg *et al*., 2015; Bazakos *et al*., 2017). The biparental mapping population used here shows that there are many QTLs with small effect sizes. As genetic variation for photosynthesis can occur in thousands of genes, all with relatively small effect sizes (Theeuwen *et al*., 2022*b*), mapping populations have more power to detect these. Therefore, to reveal small effect size genetic variation to improve photosynthesis in crops, biparental populations or multiparental populations, such as MAGIC populations, will be better suited. Genomic selection procedures can then be used to bring together all of the desirable genetic variants (Furbank *et al*., 2019; Bezouw *et al*., 2019; Morales *et al*., 2020; Theeuwen *et al*., 2022*b*).

### Identifying the causal alleles and physiological differences

QTL mapping of photosynthetic phenotypes can reveal genetic variation that can directly be used in breeding programs. However, identification of the causal genes underlying a QTL can generate additional insights on the impact of natural genetic variation on the photosynthetic functioning (Theeuwen *et al*., 2022*b*). In both *A. thaliana* and crops species a range of previous studies have revealed QTLs for photosynthesis, but hardly any of these QTLs have been used to identify the underlying causal gene(s) (Poormohammad Kiani *et al*., 2008; Jung and Niyogi, 2009; Wang *et al*., 2017; Oakley *et al*., 2018*a*; Rungrat *et al*., 2019; Goto *et al*., 2021). In this study, the QTL contributing the most to the difference between the Ely and Col nucleotypes in high to low-light transitions is QTL-2^18,500^ (Figures 2, 5 and 6). Fine mapping and quantitative allelic complementation studies revealed allelic variation in *cpFtsY* to be underlying QTL-2^18,500^. The protein encoded by *cpFtsY* is part of the chloroplast signal recognition pathway. The chloroplast signal recognition pathway regulates the posttranslational guidance of light-harvesting chlorophyll binding proteins (LHCPs) to incorporated them into the thylakoid membrane (Tzvetkova-Chevolleau *et al*., 2007). In this pathway the cpFtsY is the receptor protein that aids the LHCPs to be incorporated into the thylakoid membrane (Tzvetkova-Chevolleau *et al*., 2007).

To understand how the Ely allele of *cpFtsY* causes the differences observed in NPQ compared to the Col allele, we further discuss the physiological effects of the Ely allele. The *PsbA* gene in the Ely wild type accessions causes a reduction in NPQ, primarily caused by an increase of the basal dissipation mechanism *Φ*_NO_. This suggests passive dissipation of heat as a result of the reduced ability to accept electrons by the *PsbA* encoded protein D1. Knock-out mutants of *cpFtsY* show an impaired D1 protein repair cycle, which results in reduced photosynthetic capacity (Walter *et al*., 2015). In this study we show that the Ely allele of *cpFtsY* conveys higher NPQ when *Φ*_PSII_ is lower. This higher NPQ is caused by a higher flux of NPQ (*Φ*_NPQ_), rather than the flux of the basal dissipation mechanism *Φ*_NO_. This implies that the difference in NPQ caused by the Ely allele of *cpFtsY* is caused by an active NPQ mechanism. As a reduced PSII repair cycle via a knock-out of *cpFtsY* is not expected to cause an upregulation of an active NPQ mechanism, the Ely *cpFtsY* allele likely has an altered functionality, potentially facilitating the formation of more quenching sites. The absence of an expression difference between the Col and Ely alleles of *cpFtsY*, but the presence of a Pro-27-Ser amino acid change in the Ely allele can explain such difference. Further studies should reveal how the Ely allele of *cpFtsY* results in increased *Φ*_NPQ_.

It remains to be tested whether the Ely allele of *cpFtsY* causes the faster NPQ relaxation observed between the Col and Ely nucleotype when the PSII efficiency is reduced. Faster initiation and relaxation of NPQ is considered a desirable trait, as NPQ prevents harmful damage to the reaction centra (Rutherford *et al*., 2012). At a given *Φ*_PSII_ the Ely allele of *cpFtsY* causes increased NPQ, and when NPQ is increased 1 – q_P_ is substantially reduced (Figures 5D and 9A). The lower 1 – q_P_ indicated that the QA pool is more oxidized, which suggests that there is less chlorophyll triplets being formed and less damage to the reaction centra (Murata *et al*., 2007). Overall this means that the *cpFtsY* allele of Ely results in less photodamage, when PSII is less efficient. Having more oxidized Q_A_ also suggests the Ely allele of *cpFtsY* enables more quenching complexes (i.e. more NPQ). To test what the impact of having more efficient NPQ is on overall plant performance, we grew NILs with the Ely *cpFtsY* allele and the Col control with different light conditions, ranging from low stable light to highly fluctuating light and in a semi-protected gauze tunnel. In all conditions the biomass was reduced as a result of the Ely introgression in Col around QTL-2^18,500^ (Figure 8). The reduction is probably caused by the Ely *cpFtsY* allele dissipating too much energy. Where the Ely nucleotype showed less NPQ at higher *Φ*_PSII_ values (Figure 2D), the Ely *cpFtsY* allele shows no difference of NPQ at higher *Φ*_PSII_ (Figure 5D). From this we conclude, that the Ely *cpFtsY* allele is overprotective, but there are likely additional QTLs within the DH population that reduce NPQ at high *Φ*_PSII_ values as observed between the parental nucleotypes.

The Ely accession was found along the British railway system as a result of its resistance to triazine herbicides, commonly sprayed in the period between 1957 and 1992 to keep the British railways free of weeds (Flood *et al*., 2016*a*). The mutation in the *PsbA* gene reduced the binding affinity with triazine in the D1 protein, but consequently reduced the photosynthetic performance (Holt, 1990). The Ely wild type might compensate for this by actively over dissipating energy through *cpFtsY* allele. A compensatory mechanism to the reduced photosynthetic performance caused by a *PsbA* mutation has been describe for *Phalaris paradoxa* (Schönfeld *et al*., 1987). Without a compensatory mechanism and in the absence of triazine being sprayed, since 1992, resistant genotypes in theory should have reduced fitness, in comparison to susceptible genotypes (Holt, 1990; Warwick, 1991). The wide-spread occurrence of Ely 20 years after spraying stopped, suggests the fecundity of Ely was hardly reduced. Triazine was sprayed only for a relatively short time, it is therefore unlikely that the compensatory mechanism evolved in Ely, or any of its ancestors, and spread during this period. Instead, it is likely that the compensatory mechanism of Ely was already present in the nuclear background in which the *PsbA* mutation occurred. While we find the Ely *cpFtsY* allele to be overprotecting in relatively relaxed, well fed, conditions might favour a more protective NPQ mechanism, for example in habitats in low nutrients and with full exposure to sunlight.

The environments in which plant species evolved are generally considered dynamic rather than static, and genetic variation for the capacity to deal with such dynamic environments may exist (Murchie *et al*., 2018). Here we showed how high-throughput phenotyping in dynamic light environments revealed many QTLs responsible for differences in NPQ and related photosynthetic phenotypes. While our population segregated for only two parental lines, it is not enough to state that finding this many QTLs for photosynthetic phenotypes is a universal phenomenon, but a related study suggested similar amount of QTLs in cold stressed plants (Oakley *et al*., 2018*a*). Repeating such experiments with multiparental populations, containing multiple segregating alleles and high statistical power, could show how general these observations are. This study also shows that identifying the causal genes generates insights in the physiological processes for which natural genetic variation exists. The study of knock-out alleles has generated most insights in photosynthetic performance (Levine, 1968; Scheller *et al*., 2001; Alonso *et al*., 2003; Rochaix, 2004), however the study of allelic variation may contribute to more physiological insights. The results here suggest more quenching complexes to be present as caused by a natural allele of *cpFtsY*, which is a phenomenon so far not attributes to this gene.

## Material and method

### Plant material

In this project, we used *A. thaliana* accession Col-0 (CS76113) and Ely (CS28631). The cybrids Col^Col^, Col^Ely^, Ely^Col^ and Ely^Ely^ were previously produced by (Flood & Theeuwen *et al*., 2020). Seeds of Hun-0 were collected at the platform of the train station at Huntly, Scotland (57.444373, -2.775414) in May 2014. As haploid inducer line we used the original Col-0 *GFP-tailswap* haploid inducer, which expresses a green fluorescent protein (GFP)-tagged CENTROMERE HISTONE 3 protein in a *cenh3/htr12* mutant background (Ravi and Chan, 2010). To generate cybrids containing the Hun-0 plasmotype, a Hun-0 haploid inducer line was generated by crossing the original *GFP-tailswap* haploid inducer as a male to Hun-0 accession. Diploid F_1_ lines were selfed and amongst the F_2_ progeny, plants were selected that were homozygous for the *cenh3/htr12* mutation and carrying the *GFP-tailswap*. These were selected by vapor sterilizing the seeds, obtained from the F_1_, by exposing the seeds in a closed desiccator jar for 3 hours together with a beaker with 100 mL of bleach to which 3 ml HCl was added. Seeds were sown on ½ MS agarose plates, exposed to a minimum of 4 days of 4°C after which the plates were placed for 4 days at 25°C. Seedlings showing the typical stunted root phenotype, as described by (Wijnker *et al*., 2014), were sown on 4 x 4 x 4 cm rockwool blocks in climate-controlled chambers set at 12 hours daylight, 200 µmol/m^2^/s light, 20°C/18°C day and night temperature and 70% humidity. Plants showing the typical curled leaves and reduced fertility were selected as novel *GFP-tailswap* haploid inducers, carrying the Hun-0 plasmotype. Novel *GFP-tailswap* haploid inducer lines were confirmed by PCR genotyping using a dCAPS assay (Ravi *et al*., 2014). Consecutively, Col^Hun^, Ely^Hun^, Hun^Hun^, Ely^Hun^ and Col^Hun^ cybrids were made as described by (Flood & Theeuwen *et al*., 2020).

The doubled haploid (DH) population was made between Col-0 and Ely. F_1_ plants resulting from Col-0 X Ely were made and genotyped using Kompetitive Allele Specific PCR (KASP). The F_1_ was crossed onto the *GFP-tailswap* carrying the Col plasmotype. The resulting seeds were sown on 4 x 4 x 4 cm rockwool blocks, having the feed solution (Supplementary Table 2), with on each corner of the block one seedling. At 10 days after sowing (DAS) haploid selection started, were haploids were selected upon their smaller and narrower leaves, symmetry of rosette and overall smaller size of the plant. These potential haploids were transplanted into 7 x 7 x 7 cm pots with potting soil. Another round of selection was done at flowering stage, were haploids showed smaller flowers and a high level of sterility. The haploid lines self-fertilized producing the doubled haploid lines. The doubled haploid lines are numbered from 1 to 523, and referred to as DH1 to DH523.

For the fine mapping populations and near isogenic line (NIL) construction of QTL-2^18,500^ and QTL-4^0.25^, three independent crossing schemes were followed. To produce a NIL and/or recombinants for finemapping, for QTL-2^18,500^, DH6 and Col^Col^ were reciprocally crossed, and genotyped with KASP markers. Four F_1_s were selfed to produce 500 F_2_ plants, that were genotyped with KASP (see Genotyping fine mapping and NIL genotypes). Six of these F_2_ plants were fully homozygous for Col with a heterozygous Ely introgression around QTL-2^18,500^. Selfing two F2 plants resulted in 2497 plants, of which 523 were homozygous NILs and 57 had a heterozygous recombination within a 250-kbp region around QTL-2^18,500^. Each of these 57 recombinants was selfed and the homozygous recombinant was selected, and subsequently used for phenotyping. As QTL-4^0.25^ is located close to the *FRIGIDA* (*FRI*) gene, known to convey many pleiotropic effects, and possibly the observed *Φ*_PSII_ phenotype, one NIL was made including the *fri* allele of Col and one including the *FRI* allele of Ely. For the NIL including the *fri* allele of Col, Col^Col^ was crossed to DH9, and subsequently propagated by selfing until in the F_4_ a NIL was selected. For the NIL including the *FRI* allele of Ely, Col-0 was crossed to DH466. Then the F_1_ was backcrossed to Col-0, for two generations, and the BC2 was selfed, to obtain 3 independent NILs.

For quantitatively testing whether candidate genes are the causal genes for the two QTLs, allelic complementation tests and different RIL were used. For the RILs, we used the F9 generation of 163 RILs from Bur-0 X Col-0, 164 RILs from Can-0 x Col-0 and 163 RILs from Shah x Col-0, obtained from the Versailles Arabidopsis Stock Centre and described by (Simon *et al*., 2008). For the allelic complementation experiment, SALK T-DNA insertion lines *cpftsy* (N658281) and *kea2* (N677716) were obtained from the Nottingham Arabidopsis Stock Centre. These lines were genotyped and confirmed to be homozygous for the T-DNA insertion. The seeds of Col-FRI and Col-FRI-flc3 were obtain from the Max Planck Institute for Plant Breeding Research, Cologne.

### Genotyping DH population

Genotyping of the DH population was done by extracting DNA via CTAB extraction and consecutively the Hackflex protocol was used to do the library preparation (Gaio *et al*., 2022). Samples were pooled, and the fraction showing reads of roughly 500 bp in size were selected and sequenced for on average 1X whole genome coverage sequencing with Novogene (UK) Ltd. The SNP and indel calling workflow consists of four steps: (1) read trimming, (2) read alignment, and (3) variant calling. Step 1: Reads were trimmed using Cutadapt (Martin, 2011) (version 1.18). This step clipped sequences that matched at least 90% of the total length of one of the adapter sequences provided in the NEBNext Multiplex Oligos for Illumina (Index Primers Set 1). In addition, it trimmed bases from the 5’ and 3’ ends of reads if they had a phred score of 20 or lower. Reads shorter than 70 bp after trimming were discarded. Step 2: Trimmed reads were aligned to a modified version of the *A. thaliana* Col-0 reference genome (TAIR10, European Nucleotide Accession number: GCA_000001735.2) which contained an improved assembly of the mitochondrial sequence (Sequence Read Archive accession number: BK010421) (Sloan *et al*., 2018) using bwa mem (version 0.7.10-r789) (Li, 2013) with default parameters. The resulting alignment files were sorted and indexed using samtools (version 1.3.1) (Li *et al*., 2009). Alignment files of libraries generated from the same accession were merged using Picard MarkDuplicates (https://broadinstitute.github.io/picard/), called through the GATK suite (version 4.0.2.1) (McKenna *et al*., 2010). Picard MarkDuplicates was also used to mark duplicate read pairs, using an optical duplicate pixel distance of 2500, which is appropriate when working with patterned Illumina flowcells. Step 3: SNPs and indels were called by running FreeBayes (Garrison and Marth, 2012) (version 1.3.1-dirty) with alignment files of all samples as input, using default parameters.

After the filtering steps, the read coverage came to an average of 1.2X per DH genotype with a relatively narrow distribution around the mean (Supplementary Figure 4A). With the low coverage variant calling data per DH genotype, individual variants could not reliably be used as genotyping markers. Instead a custom pipeline was built to call genotypes on the basis of different sizes of windows. Several filtering steps were used to remove very low quality variants, that represent potential false positive calls, and optimize the window genotype calling. Firstly only the two best, i.e. most common, alleles were considered for each given variant, as only two alleles should segregate in a biparental population. Variants with read coverage across all samples (DP) below 50 and above 750 were excluded, also variants with a quality (QUAL) lower than 10 were removed and quality, and normalized for the number of observed alternative allele (QUAL/AO), lower than 10 were removed. We determined the reference and alternative allele count to calculate the minor allele count and minor allele frequency, and excluding missing calls for a genotype. Next variants that are deviant of the expected segregation, but allowing variants showing segregation distortions, were excluded. For this we used a custom R script that calculated for every variant the average minor allele count of the 100-kbp region around the given variant, and used a binomial distribution to determine whether it falls within the 95% confidence interval. If the minor allele count of the variant is outside the 95% confidence interval of the 100-kbp region around it, that variant is excluded from the analysis. As a last step for the window sizes of 10, 25, 50, 100 and 250 kbp the genotype was determined using a custom R script. A window was genotyped as the reference allele if more than 50% of the variants was called as reference, and likewise the window was genotyped as alternative allele if more than 50% was called as alternative. Windows with no variants or less than 5% of the average number of variant for a given DH genotype, were left undetermined (NA). The resulting genotype files were used in all downstream analyses.

Downstream filtering of DH genotypes and markers was done by custom R scripts. DH genotypes were removed with not enough variants leaving 485 DH genotypes and the parental genotypes. Next, DH genotypes representing extremely high cross overs numbers, an indication of DNA contamination or outcrossing, were excluded. The genotyping data with the 250-kbp window size was generated based on the most variants and thus most accurate genotype call, without getting too big to miss large numbers of crossovers. This resulted in on average 8.2 cross overs per DH genotype, with 13 DH genotypes showing more than 25 cross overs, in patterns representative of DNA contamination or outcrossing. Removing these 13 DH genotypes left 472 DH genotypes. Next, markers were excluded with too high cross over rates over the entire DH population, representative of windows with difficult-to-call genotypes. Windows with more than 40 apparent cross-overs within the whole population, all showing no linkage disequilibrium with adjacent windows, were excluded for downstream analyses, which reduced the number of markers from 461 to 456. These 456 markers represent the 250-Kbp windows. In the resulting population, the average cross over per DH genotype per chromosome is 1.29 (Supplementary Figure 4B). Across the genome, relatively many cross overs were observed on the top of chromosomes 1, 4 and 5 (Supplementary Figure 4C), a finding in line with increased cross over ratio on the ends of chromosomes primarily observed on the male side of meiosis (Giraut *et al*., 2011).

Furthermore, duplicate DH genotypes, either due to sampling mistakes, DNA contaminants or seed contaminants, were excluded. Resulting in 449 DH genotypes and the two parental genotypes. Using the 250-kbp window genotype dataset, the genetic map was constructed to account for potentially larger structural variation and used for later QTL mapping approaches. For this, the Rqtl2 package was used (Broman *et al*., 2003)(version 1.47-9). Default filtering steps were performed. Then the genetic map was constructed using the Kosambi mapping function, and the marker order as given by the physical position (Supplementary Figure 4A). The genetic map size per chromosome ranged between 121.8 cM for chromosome 1 and 75.4 cM for chromosome 4 in line with the most up to date genetic map of *A. thaliana* (Meinke *et al*., 2009). Segregation distortion was analyzed and found to show a distortion on chromosome 2, physical mapping position 16.250 Mbp. Segregation distortions are not uncommon in *A. thaliana* mapping populations (Salomé *et al*., 2011), and could be a sign of seed dormancy or lethal epistatic interactions. In this population there might be a bias introduced as a result of haploid selection, as this was done on the typical narrow leaves of a haploid, but this trait could also have segregated in this population. Despite the segregation distortion, the DH population is large enough to allow for QTL mapping, even in the region showing segregation distortion.

### Genotyping fine mapping and NIL genotypes

For all genotyping steps in this project, DNA extraction was done in 96 deep well plates. Single leaves were harvested from individual plants and placed in the deep well plates, snap frozen in liquid nitrogen and ground with a Retsch MM300 TissueLyser. 100 mL extraction buffer (200 mM Tris-HCl, 25 mM EDTA and 1% SDS) was prepared by adding 40 µL of 20 mg/mL RNase A. 500 µL of extraction buffer including RNase A was added to each well, and incubated at 37 ^°^C for 1 hour, and inverted every 15 minutes. To pellet the debris the plates were centrifuged for 5 minutes at 3000 x g. In a new deep well plate 130 µL Kac buffer (98.14 g KAc and 3.5 mL Tween were added to 160 mL H_2_O and H_2_O was added to reach 200 mL) and 400 µL lysate were added. The plates were sealed and inverted for 2 minutes, and incubated on ice for 10 minutes. To pellet the debris the plates were centrifuged for 5 minutes at 3000 x g, and 400 µL of supernatant was transferred to a new plate containing 440 µL Sera-Mag Speedbeads (Cytiva Europe) diluted in PEG buffer. Plates ware sealed and place on a shaking table for 30 minutes. Next, the plates were placed for 5 minutes on a magnet and the supernatant was removed and washed with 500 µL 80% EtOH, and repeated three times. The plates were left to evaporate in the fume hood for 1 hour and the DNA was resuspended in 50 µL milliQ.

Genotyping of the fine mapping populations and NILs was done with a PCR-based Kompetitive Allele Specific PCR (KASP) (He *et al*., 2014) assay. This assay discriminated for polymorphisms between Col-0 and Ely at different positions. Primers were ordered from Biolegio BV and using a working solution of 10 µM the two forward primers and one reverse primer were mixed in a 1:1:2 ratio respectively. The PCR reaction contained 4 µL KASP Master Mix (LGC Group), 0.14 µL primer mix, 5 µL milliQ and 1 µL DNA. The PCR cycle started with 15 minute activation at 94 ^°^C, followed by 10 cycles of 20 seconds at 94 ^°^C denaturation and 1 minute annealing step at 64 ^°^C (and dropping 0.6 ^°^C per cycle), followed by 26 cycles of 20 seconds denaturation at 94 ^°^C and 1 minute annealing at 55 ^°^C. HEX and FAM fluorescence readings were taken at the end using a Bio-Rad CFX96 thermocycler. All primers that worked and have been used in this project are listed in Supplementary Data 2.

### *De novo* sequencing

As input material for the *de novo* assembly of Ely wildtype, 500 mg young leaf material was used. High-quality high molecular weight (HMW) DNA was extracted following the LeafGo protocol (Driguez et al., 2021). Using 100 ng/µL HMW DNA (Qubit dsDNA Quantitation BR Kit) short reads where eliminated with the PacBio SRE Circulomics Kit. 1 µg HMW DNA (Qubit dsDNA Quantitation BR Kit) was used to do end-prepping and nick repairing, using the NEBNext Companion Module Kit (#E7180S) followed by ligation using the Oxford Nanopore Technologies (ONT) Ligation Sequencing Kit (SQK-LSK109). Roughly 25X coverage, after base calling, was generated using an ONT Flow Cell (R9.4.1) on the MinION Mk1C. Base calling was done using the “fast basecalling” option on the MinION Mk1C. De novo assembly was created using the Flye assembler, with default settings for raw ONT reads (Kolmogorov et al., 2019). The contigs where polished in 4 subsequent rounds, using Pilon in default settings (Walker et al., 2014), with the short read Illumina data generated for Ely wildtype in (Flood & Theeuwen *et al*., 2020).

### Phenotyping and statistical analysis

#### Cybrids in the Dynamic Environment Phenotyping Instrument

The data for this analysis was obtained from Flood & Theeuwen *et al*., 2020. For this analysis we took the data from the experiment that took three days, with light conditions as shown in Figure 1A. For the analysis of differences between the Ely and Col plasmotype, only the cybrids with these two plasmotypes were included. The analysis of differences between the Ely and Col nucleotype was done with the cybrids that had these two nucleotypes. As the plants were grown randomized over the growing chamber, without blocking, the statistical analysis was done with a linear mixed model with equation:

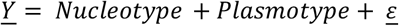

Using this equation the Best Linear Unbiased Estimates (BLUEs) were extracted over all nucleotypes, in the case of the Col versus Ely plasmotype analysis. In the case of the nucleotypes analysis, the BLUEs were extracted over all the plasmotypes. A pairwise t-test was performed using α = 0.05 with n=4.

#### NPQ relaxation experiment

This experiment consisted out of two separate experiments, one phenotyping the NPQ relaxation in darkness and one with a light intensity of 50 µmol m^2^ s^-1^. For both experiments cybrids Col^Col^ and Ely^Col^ were grown. The seeds were sown in Petri dishes on soaked filter paper with 1 mL. Then the seeds were placed in a dark cold room (4 °C) to induce vernalization. After one week these seeds were placed in a climate-controlled chamber (24 °C with a rhythm of 16/8 h day/night) for 24 h to induce germination. The germinates seeds were sown on pre-soaked rockwool supplied by Grodan B.V. (Roermond, the Netherlands) in a climate-controlled chamber. Plants were irrigated with feed solution every three days (Supplementary Table 2). The chamber had a photoperiod of 16 h with 200 µmol m^2^ s^-1^ irradiance, the temperature was 20 °C during the day and 18 °C during the night, the relative humidity was 70%. The plants were sown in a complete randomized block design, with n=10 per block.

After 14 days of growth, plants were phenotyped in the PlantScreen^TM^ system provided by Photon Systems Instruments spol. s r.o (Drásov, Czech Republic). One block of two times 10 Col^Col^ and Ely^Col^ were measured at once. All plants were dark adapted for 30 minutes, to retrieve Fo and Fm. Next plants were exposed to a sequence of 1000, 100, 1000, 100 and 1000 µmol m^-2^ s^-1^ with three minutes each. At the end of the last 1000 µmol m^-2^ s^-1^ Fm’ was calculated. Right afterwards the lights were either turned off or set to 50 µmol m^-2^ s^-1^, depending on the experiment. To avoid the influence of saturating light pulses on NPQ relaxation too much, saturating light pulses was applied to measure Fm’, with 30 s in between for a period of 300 s. Both experiments were repeated six times, and in every repetition the first measurement after the 1000 µmol m^-2^ s^-1^ was delayed by 5 s. This resulted in NPQ relaxation data every 5 s, for both cybrids, during the entire 300 s. Next, all timepoints were separately analysed using a linear model with the following model:

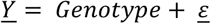

Significant differences were calculated with a threshold in the post hoc tests of α = 0.05, using the following equation.

#### Phenotyping DH population and fine mapping lines

The DH population was grown in two separate high-throughput phenotyping systems. The first system we used was the Dynamic Environmental Photosynthetic Imaging (DEPI) (Cruz *et al*., 2016), with modifications as described in Tietz *et al*., 2017. The plants were grown for 18 days in a climate-controlled chamber with a light intensity of 200 µmol m^-2^ s^-1^ with a photoperiod of 16 h. After 18 days these plants were moved to the DEPI system and acclimated for three days. At day 21 the experiment was initiated with light conditions as shown in Figure 4A. The phenotyping of all DH lines was separated over eight experiments with 224 plants in each, with n=3 or 4. Each experiment had a minimum of 14 Col^Col^ and Ely^Col^ included. Throughout the experiment growth measurements and chlorophyll fluorescence parameters (F_v_/F_m_, NPQ, NPQ_(t)_, *Φ*_PSII_, *Φ*_NPQ_, *Φ*_NO_, *q*_E_, *q*_E(t)_, *q*_I_, *q*_I(t)_, *q*_L_) were taken. The broad sense heritability was calculated using the following equation:

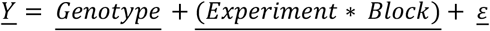

The BLUEs were calculated using a linear mixed model:

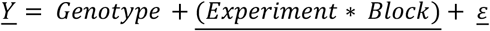

The QTL mapping was done using the MQM method in R/qtl (Broman *et al*., 2003; Arends *et al*., 2014). For this the genetic map was used with markers every 250-Kbp region. A pseudo marker was introduced every 3^th^ marker. As cross type we used selfed RIL, as DH is not supported in MQM. The positive and negative LOD scores were calculated using the effect size of the Col versus the Ely allele. The Bonferroni threshold was calculated, by correcting for the number of markers and phenotypes. In the analysis we found 69 of the 449 DH lines to possess the Cvi plasmotype instead of the Col plasmotype. Whilst conferring a phenotype effect, the analysis with and without these DH lines resulted in the same QTLs (Supplementary Figure 7). The fine mapping population was phenotyped and analysed in exactly the same way as the DH population.

The second high-throughput phenotyping system used to phenotype the DH population was the Phenovator system (Flood *et al*., 2016*b*). The DH population was germinated as described in the NPQ relaxation experiment. The plants were sown in a climate-controlled chambers, on rockwool blocks of 4×4×4 cm. They were irrigated once a week with a feed solution (Supplementary Table 3). The photoperiod was 10 h, 20/18 °C day/night, 70 % relative humidity and light intensity as shown in Supplementary Figure 10. The DH population was grown at once in the system with a complete randomized block design (n = 3). Per day during the photoperiod six measurements of *Φ*_PSII_ were taken, and nine measurements of projected leave area. Outliers were removed when either not germinated or badly established, using a R script to remove plants 1 standard deviations smaller than the average per genotype per treatment, on the basis of plant leaf area as measured during the morning of day 18 after sowing. The broad sense heritability was calculated using the following equation:

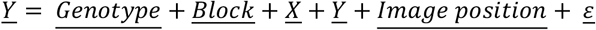

The BLUEs were calculated using a linear mixed model:

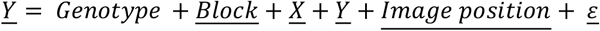

The fine mapping QTL-2^18,500^ of was done in exactly the same way as described for the analysis of the DEPI data. Phenotyping of the NILs was done in a separate experiment, with the same conditions as described above.

The three RIL populations were grown in three separate experiments in the Phenovator system. Plant were germinated as described for the Phenovator experiment performed with the DH population. Plants were grown in a climate-controlled chamber with a photoperiod of 14 h, 20/18 °C day/night, 70 % relative humidity and 200 µmol m^-2^ s^-1^. *Φ*_PSII_ was measured three times a day. The genotypes were sown in a complete randomized block design, with n = 8. All measurements were analysed using a linear mixed model approach, using the restricted maximum likelihood procedure with the lme4 package (version 1.1-30). BLUEs were calculated with the following statistical model:

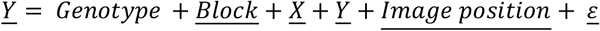

Subsequent QTL mapping was done with the MQM method as described for the DH population analysis.

#### Biomass experiments in different light regimes

For the biomass experiment cybrids Col^Col^ and Ely^Col^ and three independent NILs surrounding QTL-2^18,500^ were used. Plants were germinated as described for the NPQ relaxation experiment, and grown with a photoperiod of 12 hours. During the photoperiod the temperature was 20 °C and during the night 18 °C, and relative humidity was kept at 70%. Plants were grown on rockwool blocks and irrigated with feed solution (Supplementary Table 2) once a week. During the entire growth period the plants were grown in one climate-controlled chamber. Within the chamber three separate compartments were created, and in each compartment a different light regime was applied. The LED light systems and the controls via a ESP32 are described in (Theeuwen *et al*., 2022*a*). The three light treatments were: (1) stable light conditions with an intensity of 415 µmol m^-2^ s^-1^, (2) a sinusoidal fluctuating light regime inspired by the third day of the DEPI experiment as shown in Figure 1A and (3) a highly fluctuating light condition. The fluctuating light condition is based on measurements within a maize canopy as described in (Theeuwen *et al*., 2022*a*). Plants were grown in a complete randomized block design, with 12 blocks of 20 plants in each treatment (n = 48). The fourth treatments was grown separately in a semi-protected gauze tunnel in spring 2021. The materials and methods used for this experiment are similar to the spring experiment in 2021 as conducted for the cybrid panel as described in (Theeuwen *et al*., 2022*a*). The conditions as measured at plant level as shown in Supplementary Figure 15. The plants were sown in a complete randomized block design, with n = 40.

Plants from all four treatments were phenotyped for chlorophyll fluorescence parameters with the PlantScreen^TM^ system, using a 6 minute fluctuating light regime, yielding 37 chlorophyll fluorescence and 20 morphological parameters (Theeuwen *et al*., 2022*a*). The phenotyping was done 20 days after sowing for the plants in the climate-controlled chamber and 40 days after sowing for the semi-protected experiment. Shoot dry weight measurements for the plants in the climate-controlled chamber were taken 27 days after sowing, and 41 days after sowing for the plants in the semi-protected experiment.

Outliers were removed when either not germinated or badly established, using a R script to remove plants 1.5 standard deviations smaller than the average per genotype per treatment, on the basis of plant leaf area. Next all parameters were analysed using a linear mixed model approach, using the restricted maximum likelihood procedure with the lme4 package (version 1.1-30). Using analysis of variance with the Kenward-Roger approximation for degrees of freedom, significant differences were calculated with a threshold in the post hoc tests of α = 0.05 with Tukey correction. This revealed no differences between the three independently obtained NILs, and thus were grouped together. The analysis was done on each treatment separately, using the following equation:

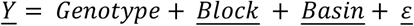

## Data availability

The raw data files with genotyping and phenotyping data will be available via Zenodo.

## Code availability

The scripts for the genotyping pipeline, the de novo assembly and QTL mapping will be available from GitLab (https://git.wur.nl/tom.theeuwen/ely-npq.git).

## Acknowledgements

The seeds of Col-FRI and Col-FRI-flc3 were kindly provided by George Coupland. We would like to thank Sofie Hofman, Max van der Sandt and Sietze Wals for their help in experimental work. Also we would like to thank Ben Auxier, Emilie Wientjes, Maarten Koornneef, René Boesten and Roel van Bezouw for discussions that help shape this work. Maarten Koornneef is also acknowledged for his feedback on the manuscript.

## Author contributions

TPJMT, JH, and MGMA conceived and designed the study. TPJMT, LLL, SP, HB, KL, JD, PJF, CH, FFMB, RW and DH performed and analysed experiments. TPJMT oversaw the whole project and DMK, JH and MGMA provided steering during the project. TPJMT wrote the manuscript, with feedback from JH and MGMA.

## Supplementary Tables

**Supplementary Table 1.**
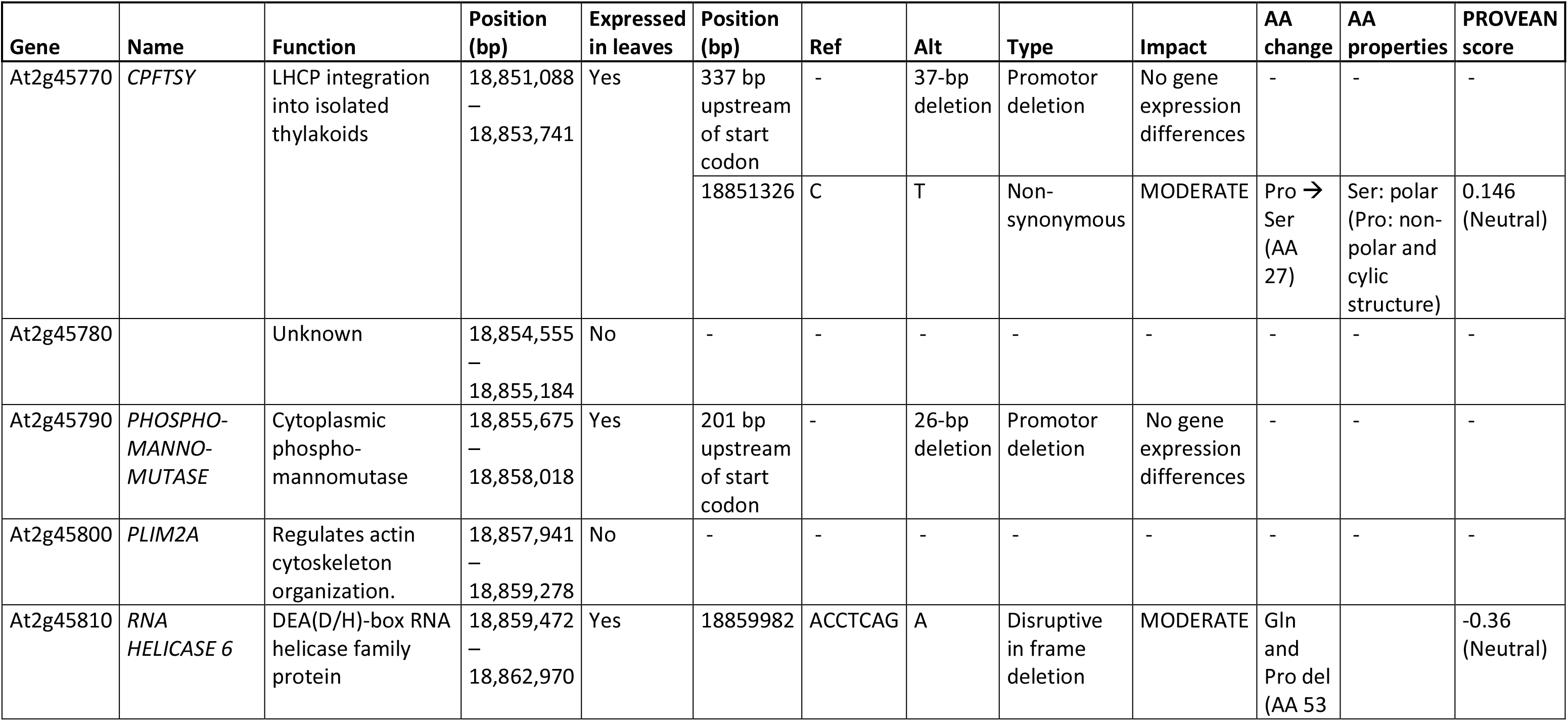
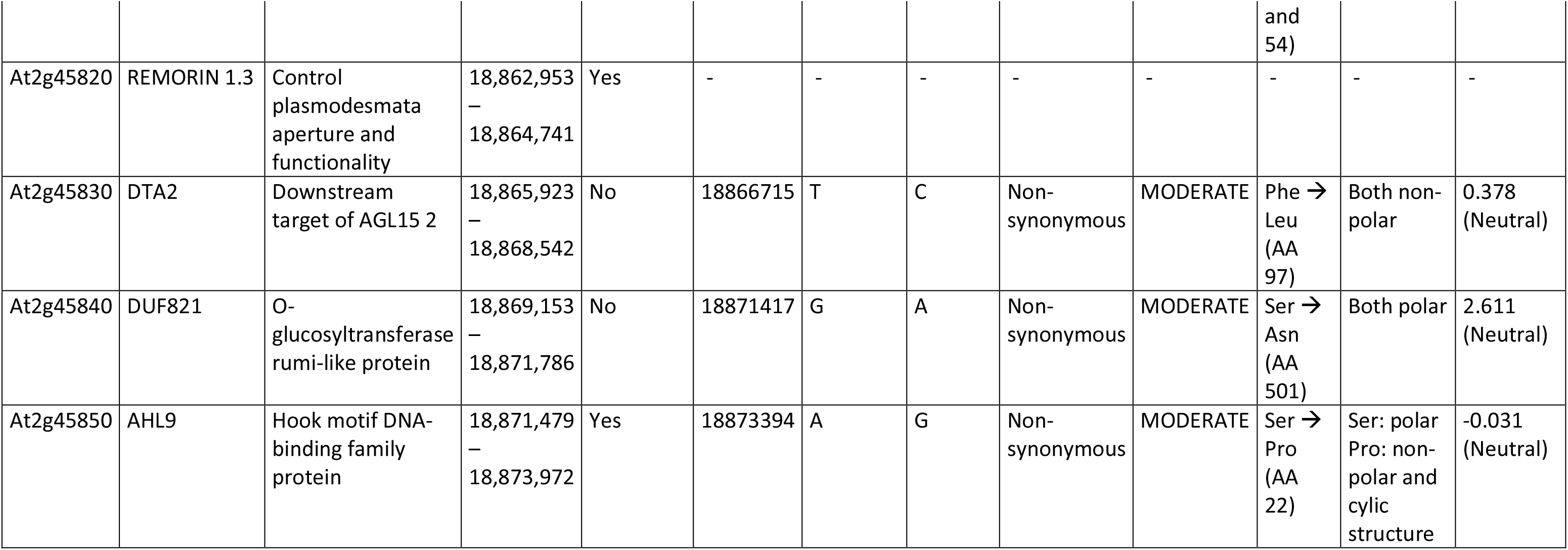
Overview of genes and genetic variation between Ely and Col in the QTL-2^18,500^. For each gene within the QTL the abbreviation, function, position (bp) and whether it is expressed in leaves, is shown. The genetic variant within or upstream of the gene are shown, with the positions, and the reference and alternative alleles. The predicted impact and amino acid change are given in the case of non-synonymous SNPs. The impact of intergenic mutations is difficult to predict, and hence excluded, with the exception of two deletions in the region that could be the promotor.

**Supplementary Table 2.** Nutrient solution as used for growing A. thaliana on rockwool substrate.

|  | Concentration | Unit |
| --- | --- | --- |
| NH<sub>4</sub> | 1.7 | mmol/L |
| K | 4.13 | mmol/L |
| Ca | 1.97 | mmol/L |
| Mg | 1.24 | mmol/L |
| NO<sub>3</sub> | 4.14 | mmol/L |
| SO<sub>4</sub> | 3.14 | mmol/L |
| P | 1.29 | mmol/L |
| Fe* | 21 | µmol/L |
| Mn | 3.4 | µmol/L |
| Zn | 4.7 | µmol/L |
| B | 14 | µmol/L |
| Cu | 6.9 | µmol/L |
| Mo | 0.5 | µmol/L |
| EC | 1.4 | mS/cm |
| pH** | 6.1 |  |

\*Consists of 50% Fe-DTPA 3% and 50% Fe- EDDHSA 3% \*\*pH adjusted with Potassium hydroxide and Sulphuric acid

## Supplementary Figures

**Supplementary Figure 1.**
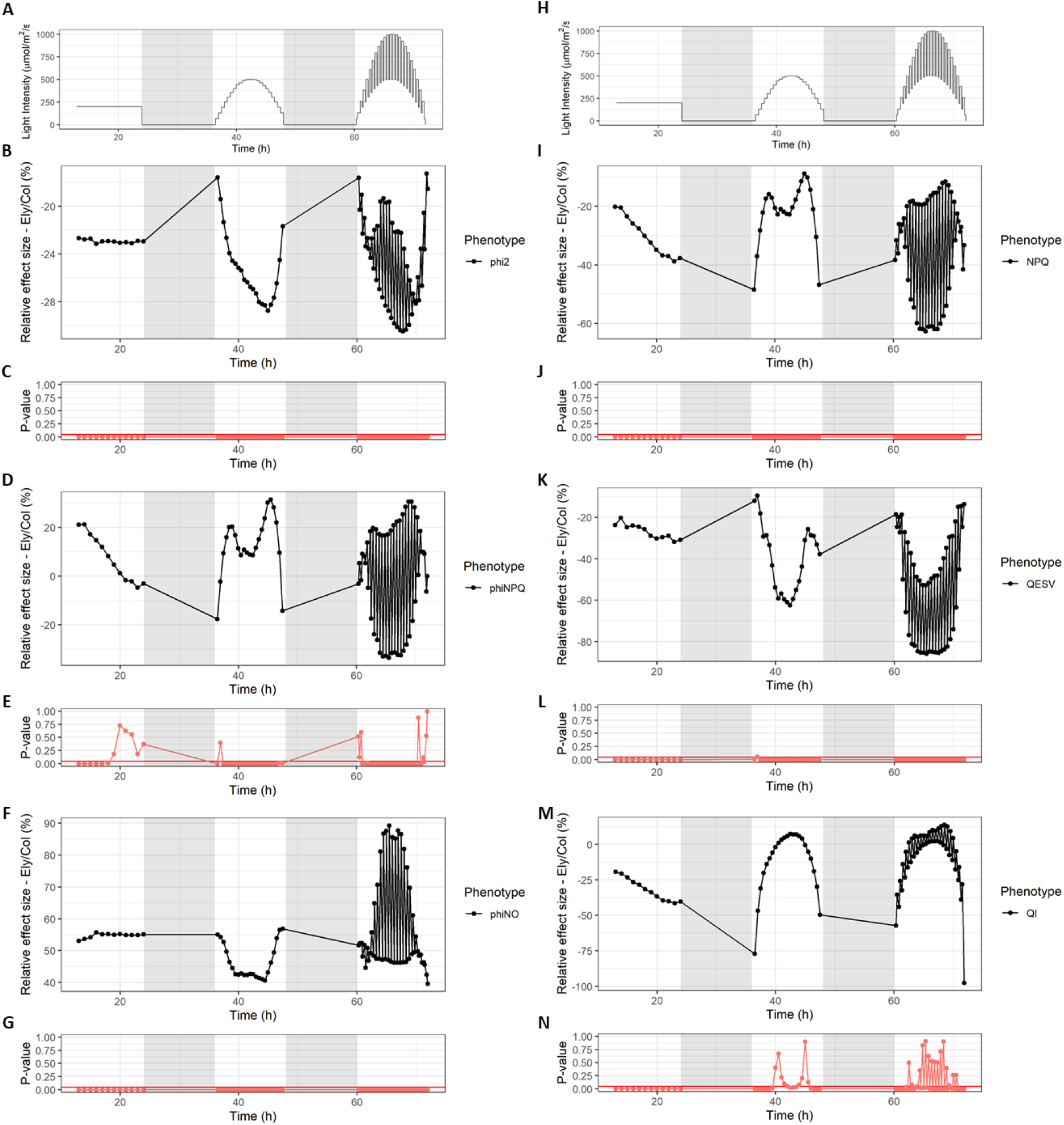
Effect size of Ely plasmotype possessing a Ser264Gly amino acid substitution for different photosynthetic parameters in dynamic light conditions. Panel A and H show the dynamic light regime to which the plants are exposed. For Φ_PSII_, Φ_NPQ_, Φ_NO_, NPQ, q_E_ and q_I_ the differences between the Col and Ely plasmotype are given, averaged over all nucleotypes. Below each panel with a phenotype, the p-value is given representing a t-test (n=4).

**Supplementary Figure 2.**
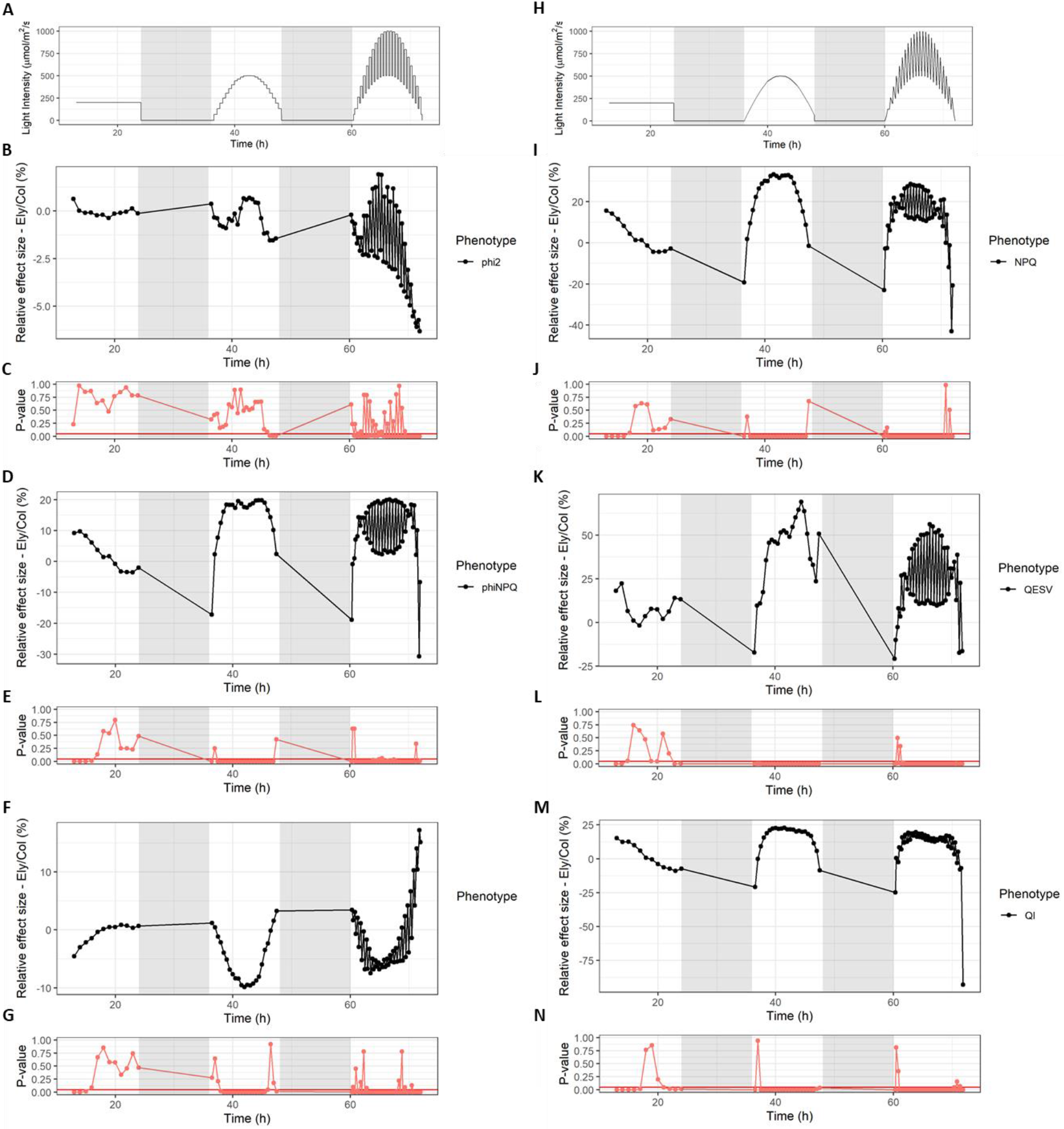
Effect size of Ely nucleotype for different photosynthetic parameters in dynamic light conditions. Panel A and H show the dynamic light regime to which the plants are exposed. For Φ_PSII_, Φ_NPQ_, Φ_NO_, NPQ, q_E_ and q_I_ the differences between the Col and Ely nucleotype are given, averaged over all plasmotypes. Below each panel with a phenotype, the p-value is given representing a t-test (n=4).

**Supplementary Figure 3.**
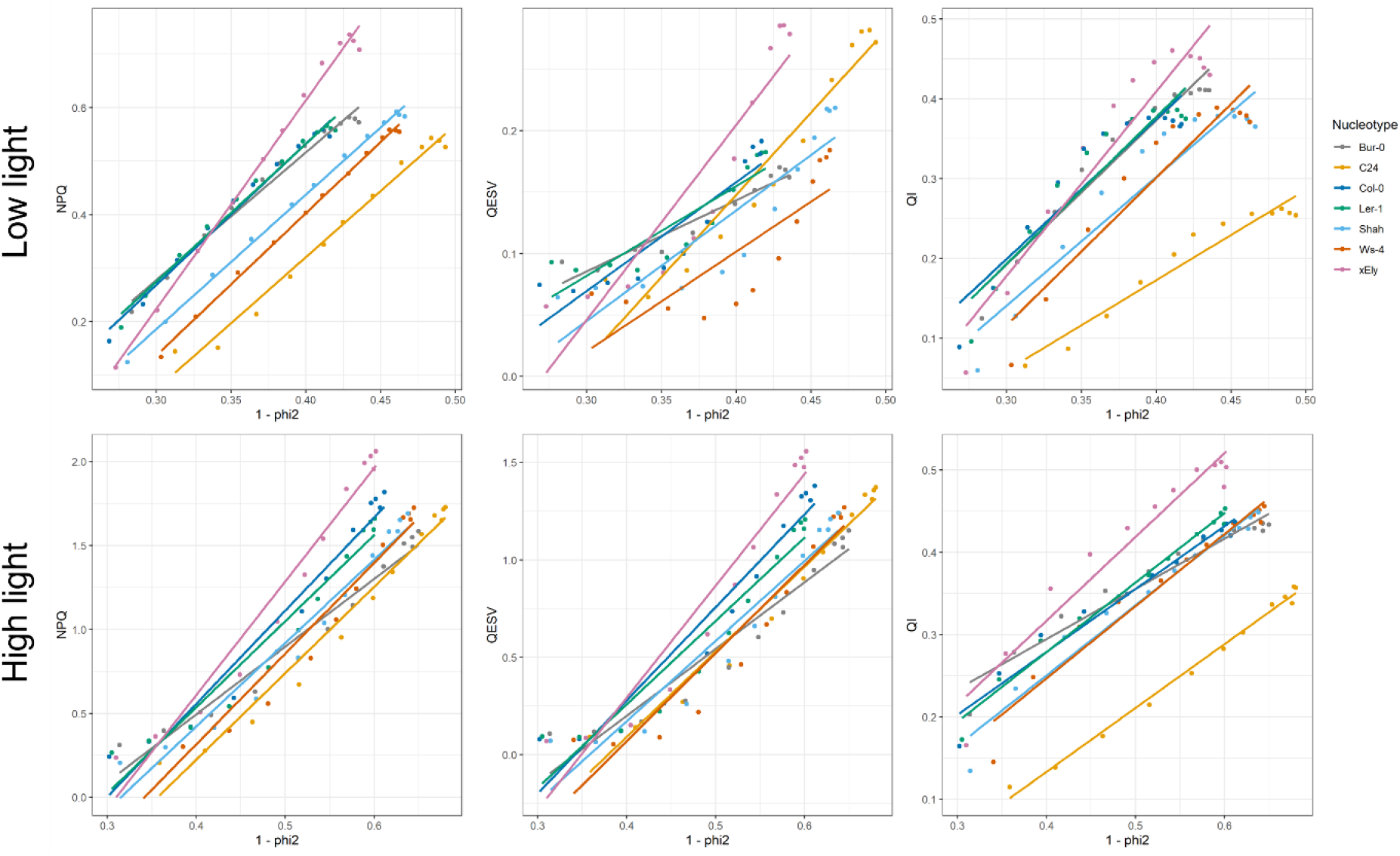
NPQ and components there off, q_E_ and q_I_, plotted against the inefficiency of PSII. The top panels visualize the response after high to low-light transitions and the bottom panels visualize the response after low to high-light transitions. In all panels the averages of the nucleotypes are given, averaged over the plasmotypes. The light intensities are taken from a fluctuating light day, as shown in Figure 2A.

**Supplementary Figure 4.**
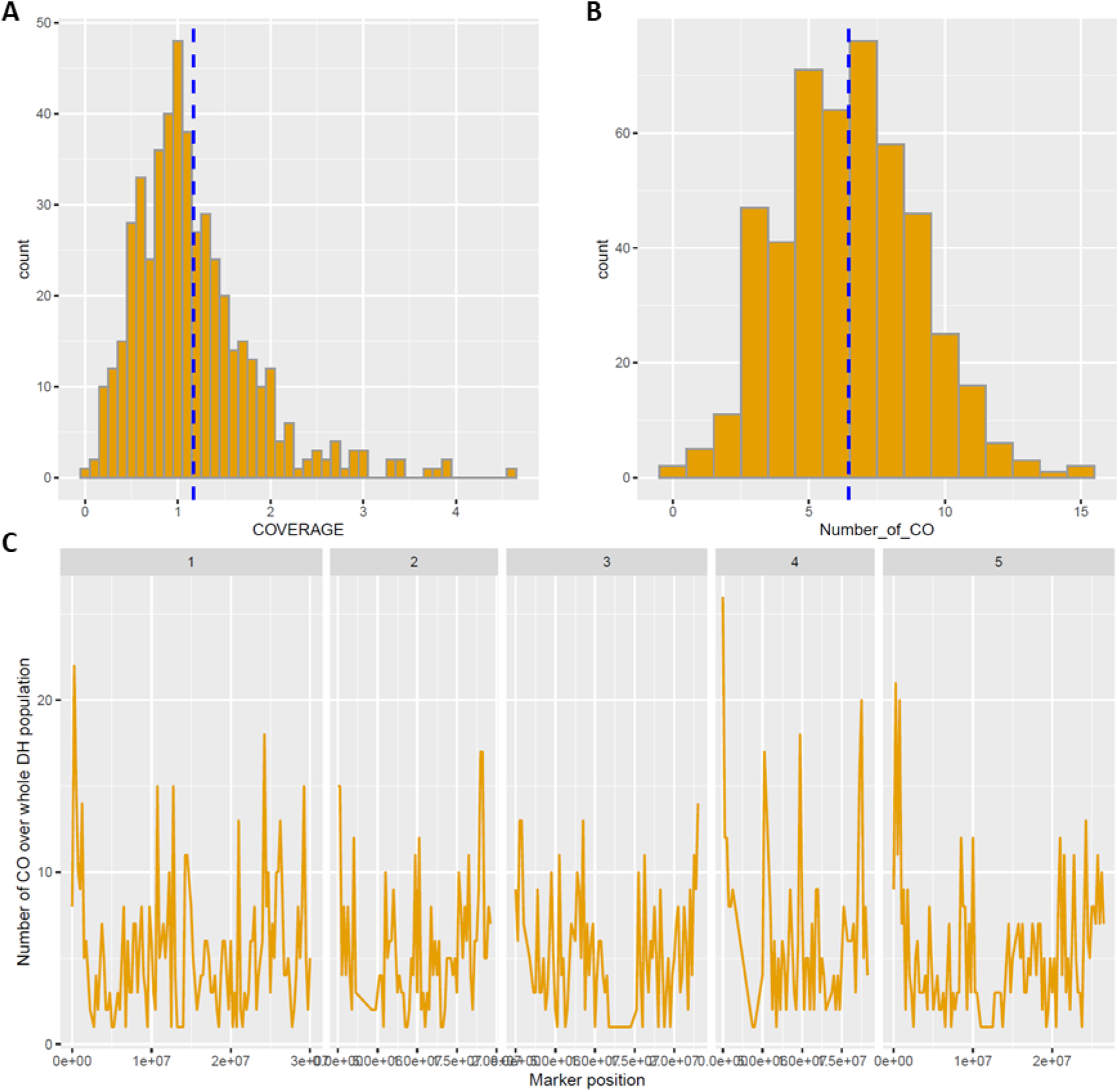
Properties of DH population. A) Distribution of read coverage per DH genotype. B) Distribution of cross overs per DH genotype. C) Average number of cross overs per 250Kbp window.

**Supplementary Figure 5.**
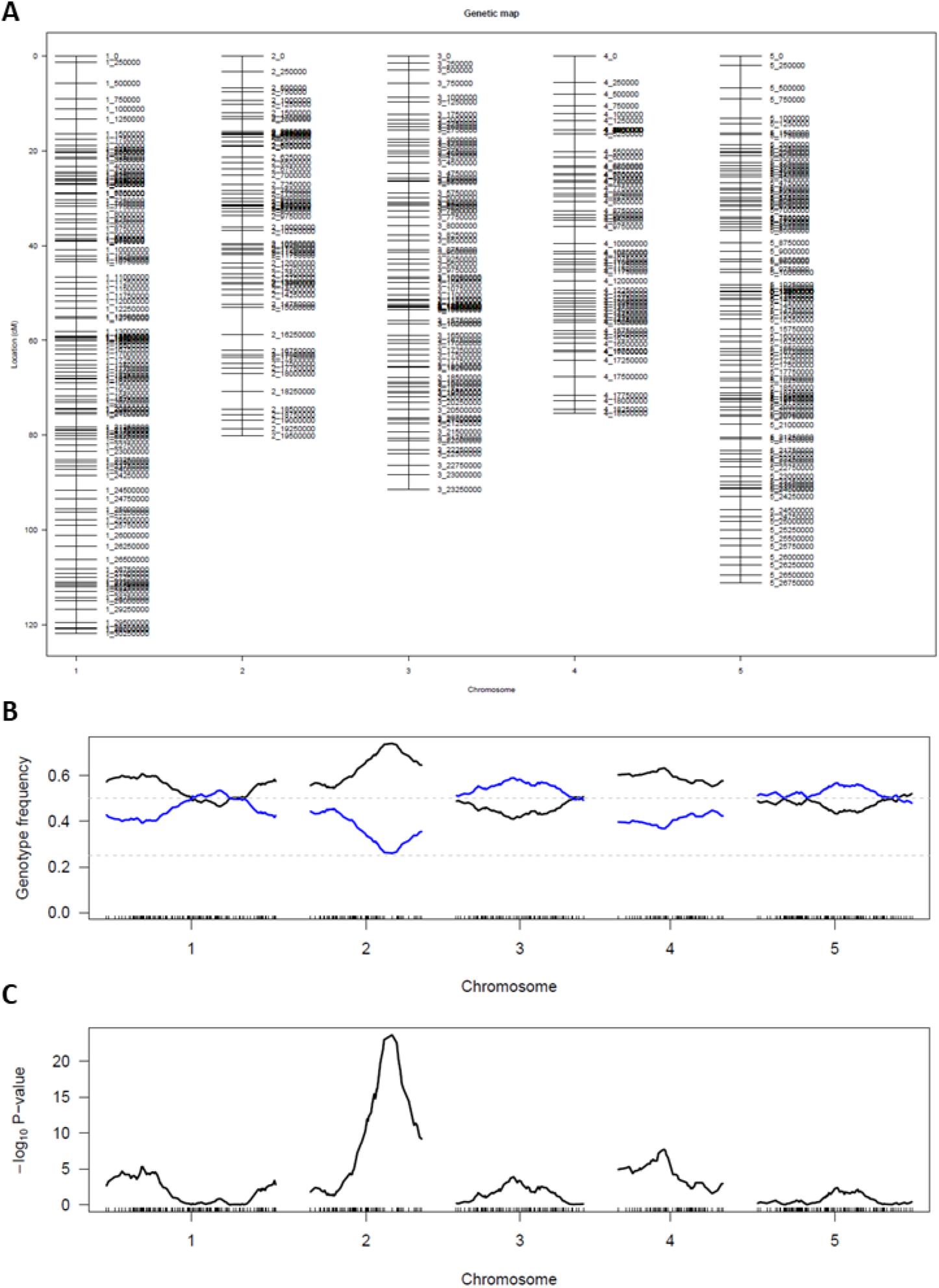
Genetic map and segregation distortion of Ely X Col DH population. A) genetic map based on markers representing 250kbp windows. B) Segregation distortion, with genotype frequency of the Ely alleles in blue and Col alleles in black. C) Significance tests for segregation distortion, with a sharp peak on chromosome 2, at physical mapping position 16.250Mbp.

**Supplementary Figure 6.**
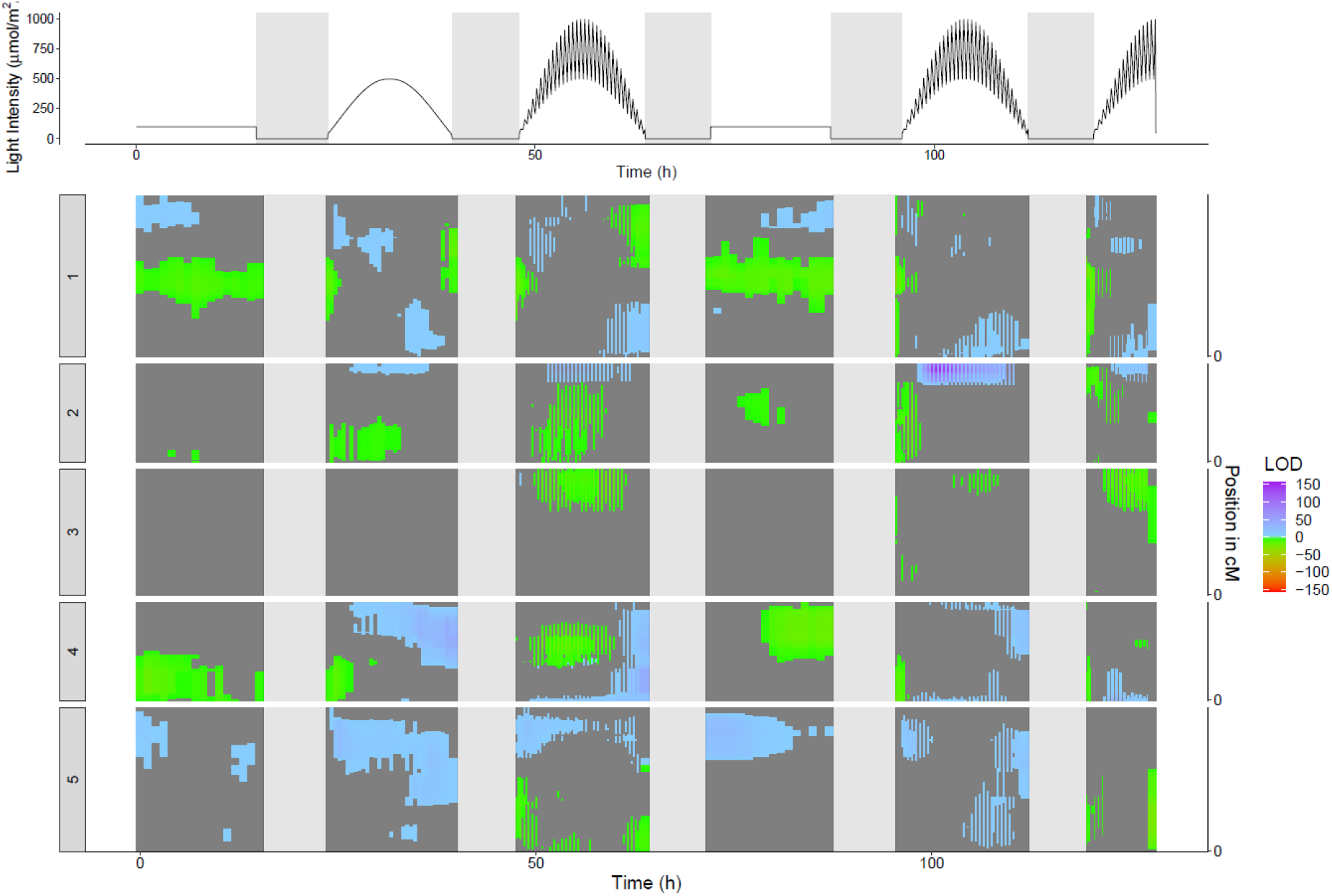
MQM plot for NPQ with DH genotypes having the Cvi plasmotype removed. This leaves 370 DH lines. In this plot the Col-0 phenotype is the control. When the Ely allele has a trait-enhancing phenotype it is indicated with a positive LOD score (blue), and when the Ely allele has a trait-reducing phenotype, it is indicated with a negative LOD score (green).

**Supplementary Figure 7.**
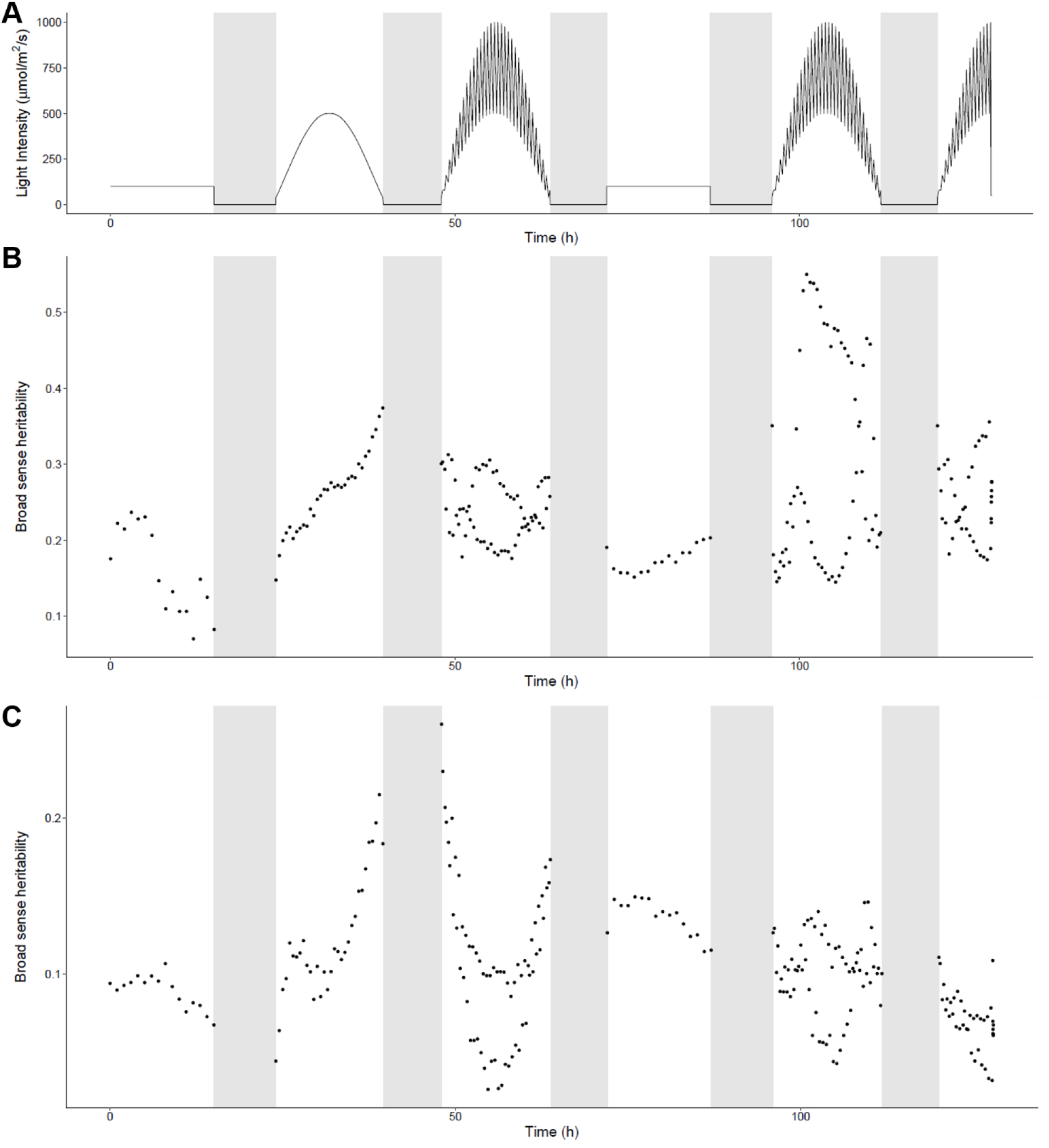
Broad sense heritability (H^2^) as observed in the DH population in the fluctuating light experiment. A) Light conditions as plants were exposed during the experiment, and matching the H^2^ in the panels below. B) H^2^ for the capacity of NPQ. C) H^2^ for the efficiency of PSII.

**Supplementary Figure 8.**
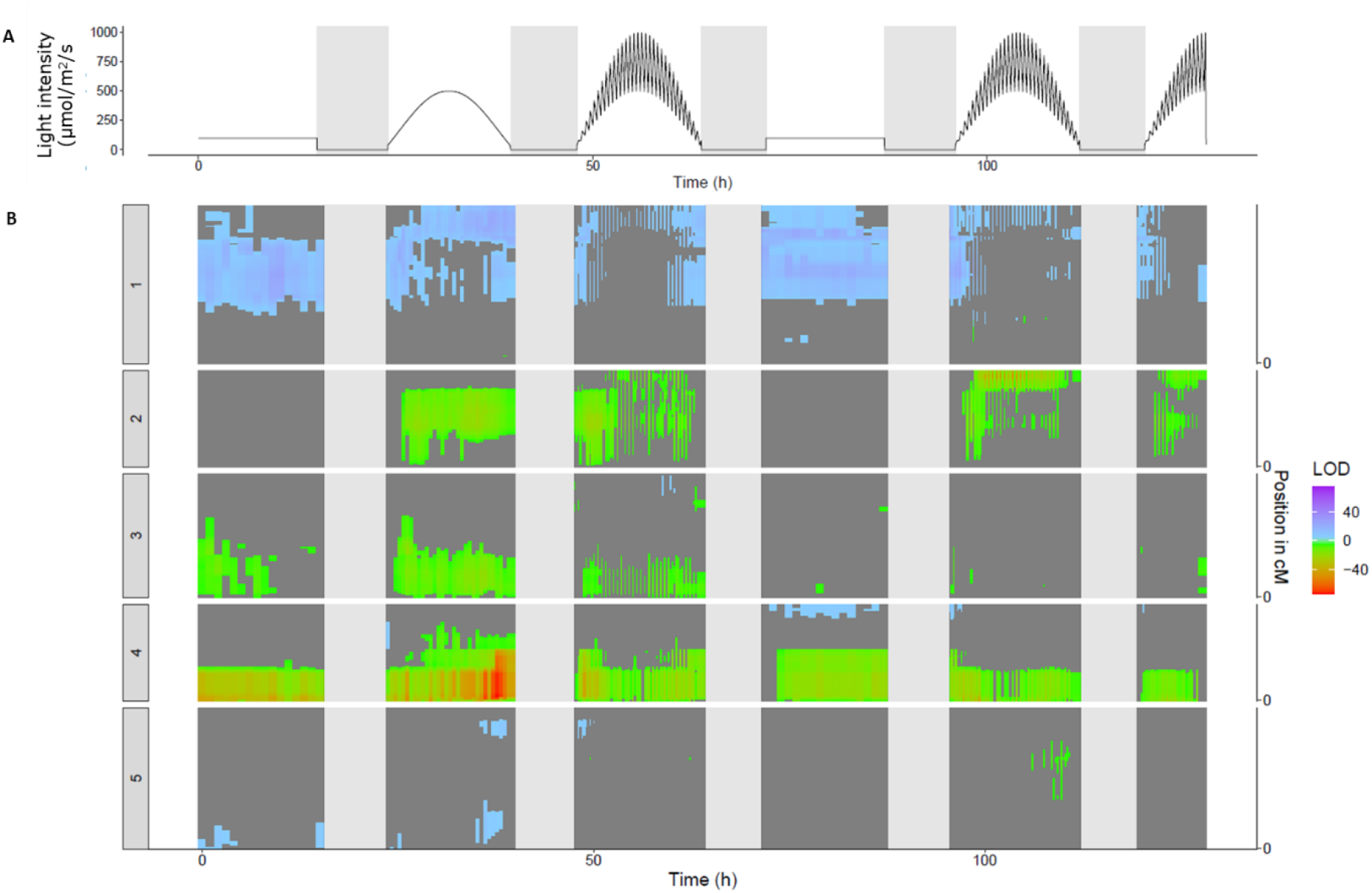
QTL map for Φ_PSII_ in DEPI. A) Represents the light intensities during the experiment, where t = 0 h is the moment when lights turned on, day 21 after sowing. In the first 21 days plants were grown at a light intensity of 200 µmol m^2^ s^-1^. B) Vertical representation of QTL mapping over time, the times match the light intensities are shown in panel A. LOD scores are represented in positive values if the effect size of the Ely allele of a given marker on that time point is higher as compared to Col allele. Negative values are given when the Ely allele induces a lower effect as compared to Col. The dark grey background indicates markers that do not pass a naive Bonferroni threshold (LOD threshold of 4.8).

**Supplementary Figure 9.**
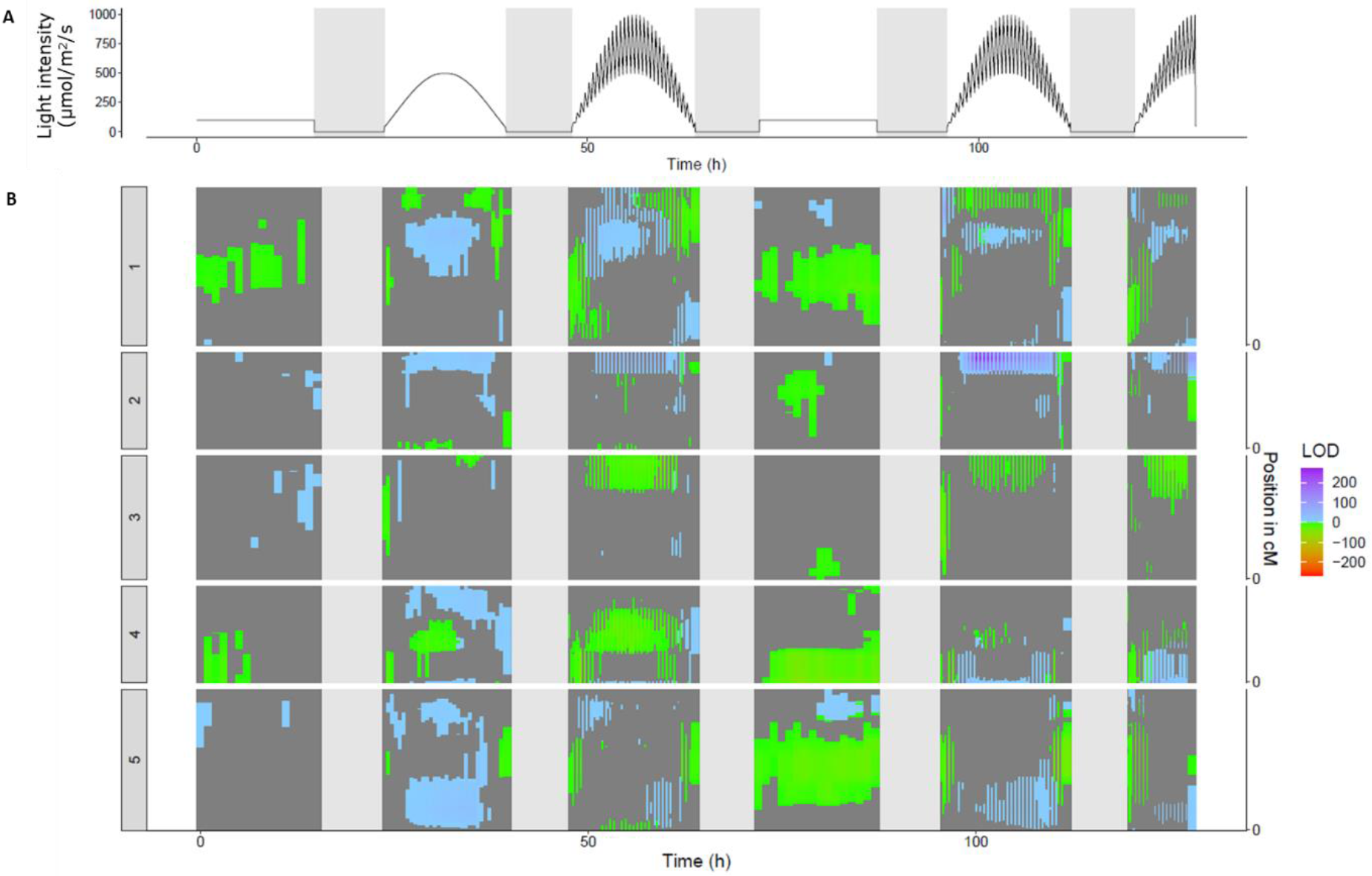
QTL map for q_E_ in DEPI. A) Represents the light intensities during the experiment, where t = 0 h is the moment when lights turned on, day 21 after sowing. In the first 21 days plants were grown at a light intensity of 200 µmol m^2^ s^-1^. B) Vertical representation of QTL mapping over time, the times match the light intensities are shown in panel A. LOD scores are represented in positive values if the effect size of the Ely allele of a given marker on that time point is higher as compared to Col allele. Negative values are given when the Ely allele induces a lower effect as compared to Col. The dark grey background indicates markers that do not pass a naive Bonferroni threshold (LOD threshold of 4.8).

**Supplementary Figure 10.**
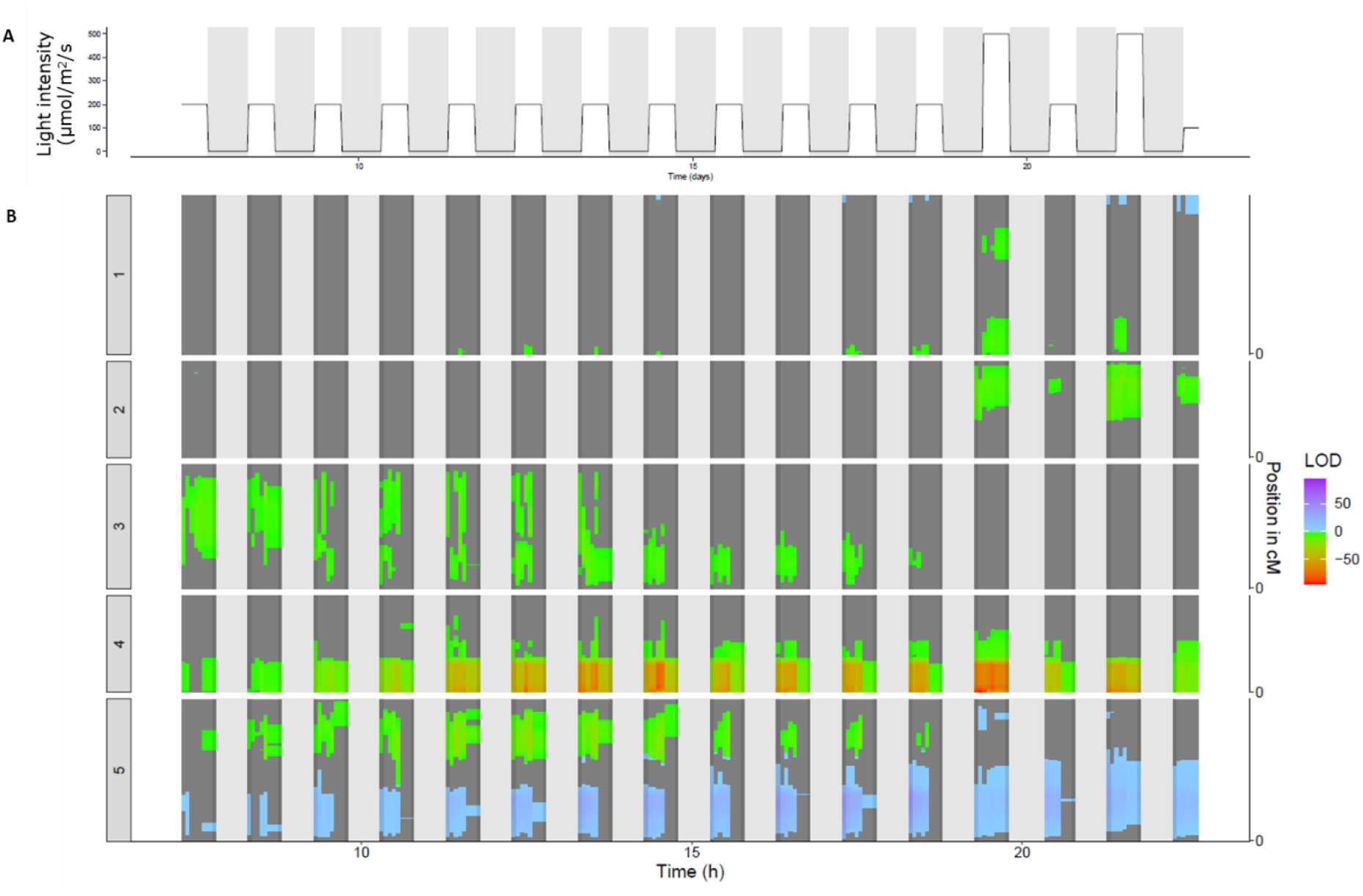
DH population in Phenovator system for Φ_PSII_. A) Represents the light intensities during the experiment, where the time is days after sowing. B) Vertical representation of QTL mapping over time, the times match the light intensities are shown in panel A. LOD scores are represented in positive values if the effect size of the Ely allele of a given marker on that time point is higher as compared to Col allele. Negative values are given when the Ely allele induces a lower effect as compared to Col. The dark grey background indicates markers that do not pass a naive Bonferroni threshold (LOD threshold of 4.8).

**Supplementary Figure 11.**
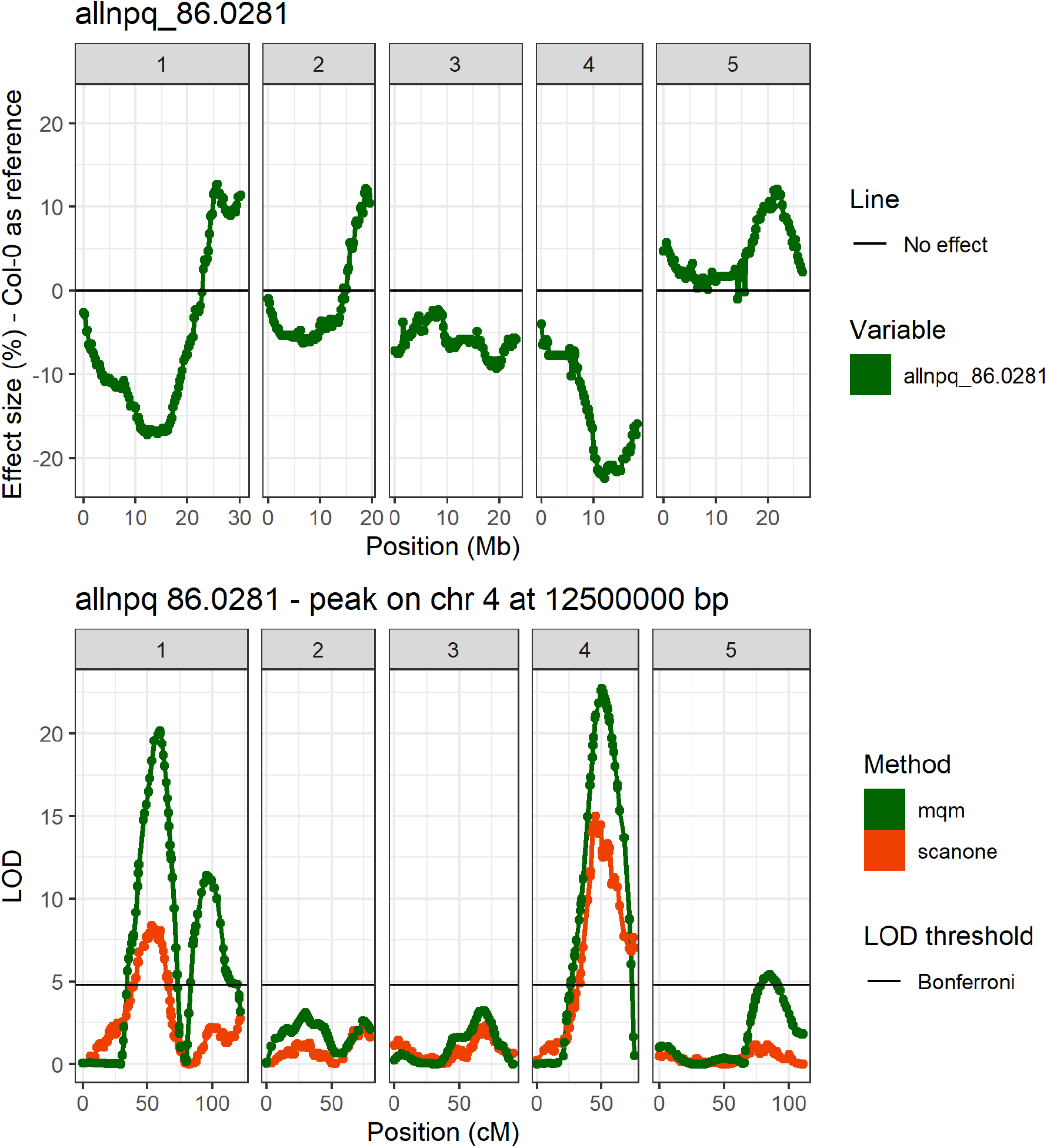
Example of a timepoint at which multiple QTLs are found for NPQ, having opposing effect sizes. A) Shows the effect size of a marker, when the Ely allele is compared to the Col allele, a positive effect size means the Ely allele causes higher NPQ. B) Shows the QTL maps at the same timepoint.

**Supplementary Figure 12.**
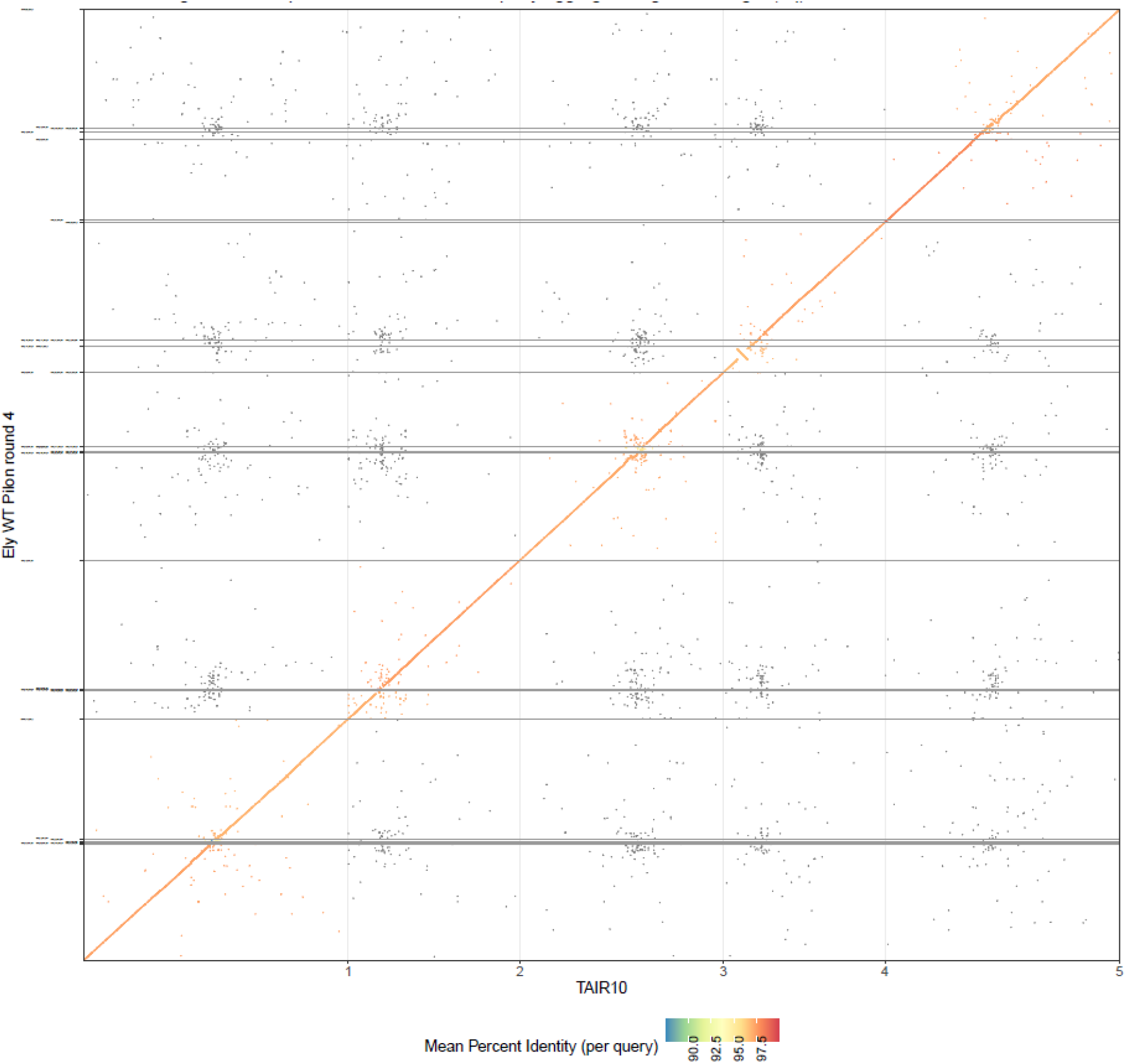
Dot plot of the de novo assembly of the Ely nuclear genome versus the Col reference genome of TAIR10.1.

**Supplementary Figure 13.**
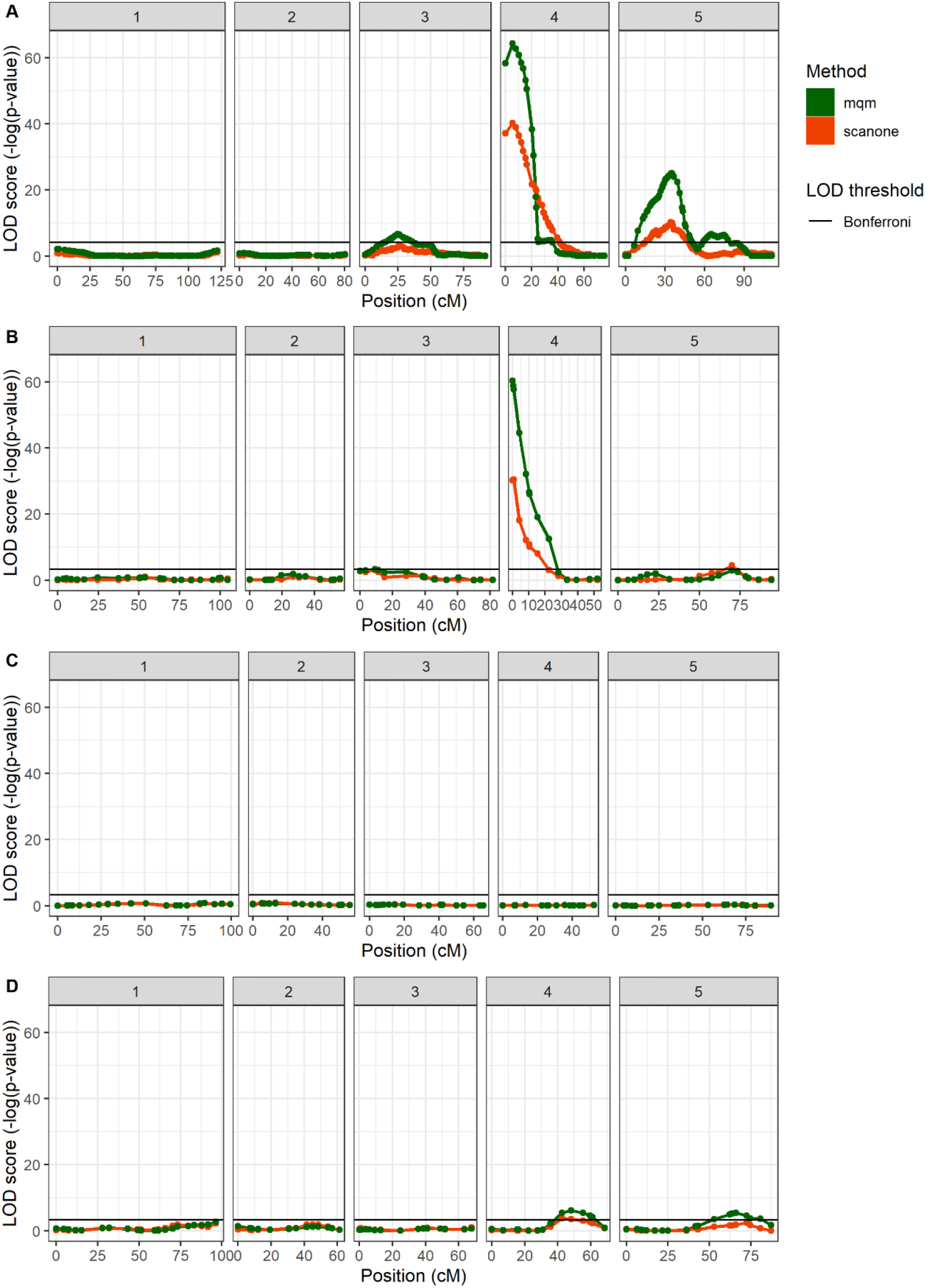
Comparison between the DH and RIL populations at 16.5 DAS for Φ_PSII_ for QTL-4^0.25^. Note how the X-axis is on a cM scale representing the genetic map of every population independently. A) Ely x Col DH QTL map, showing the highest association at 250 Kbp (with a marker every 250 Kbp). B) Can x Col RIL QTL map, showing the highest association at 651 Kbp (there are not markers to the left of this position). C) Bur x Col RIL QTL map, showing no significant association at this timepoint. D) Sha x Col RIL QTL map, showing no significant association at the beginning of chromosome 4.

**Supplementary Figure 14.**
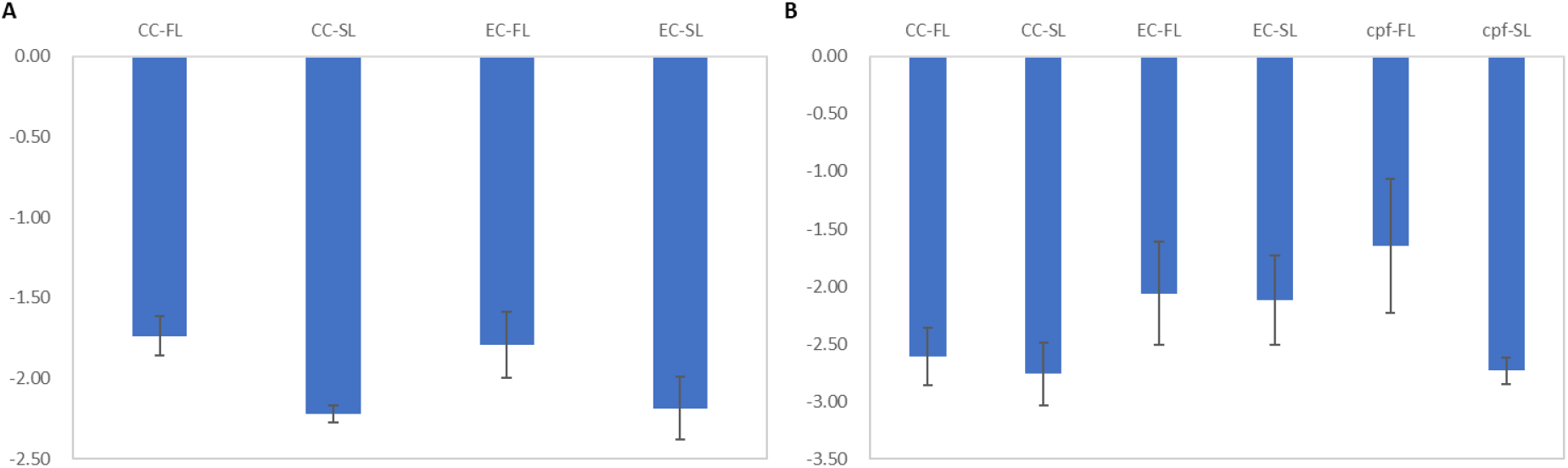
RT-qPCR results for genes in QTL-2^18,500^. A) delta-Ct values for the primer pair on PMM, with Col^Col^ and Ely^Col^ during stable light and fluctuating light (24 hours later). B) delta-Ct values for the primer pair on cpFtsY, with Col^Col^, Ely^Col^ and a cpftsy T-DNA line during stable light and fluctuating light (24 hours later). For all samples the average of five reference genes was used to calculate the delta-Ct values. These reference genes are PP2AA3, PPR, UBC9, UBQ7, SAND. All delta-Ct values are calculated with n=6. Between genotypes no significant differences are observed (α = 0.05).

**Supplementary Figure 15.**
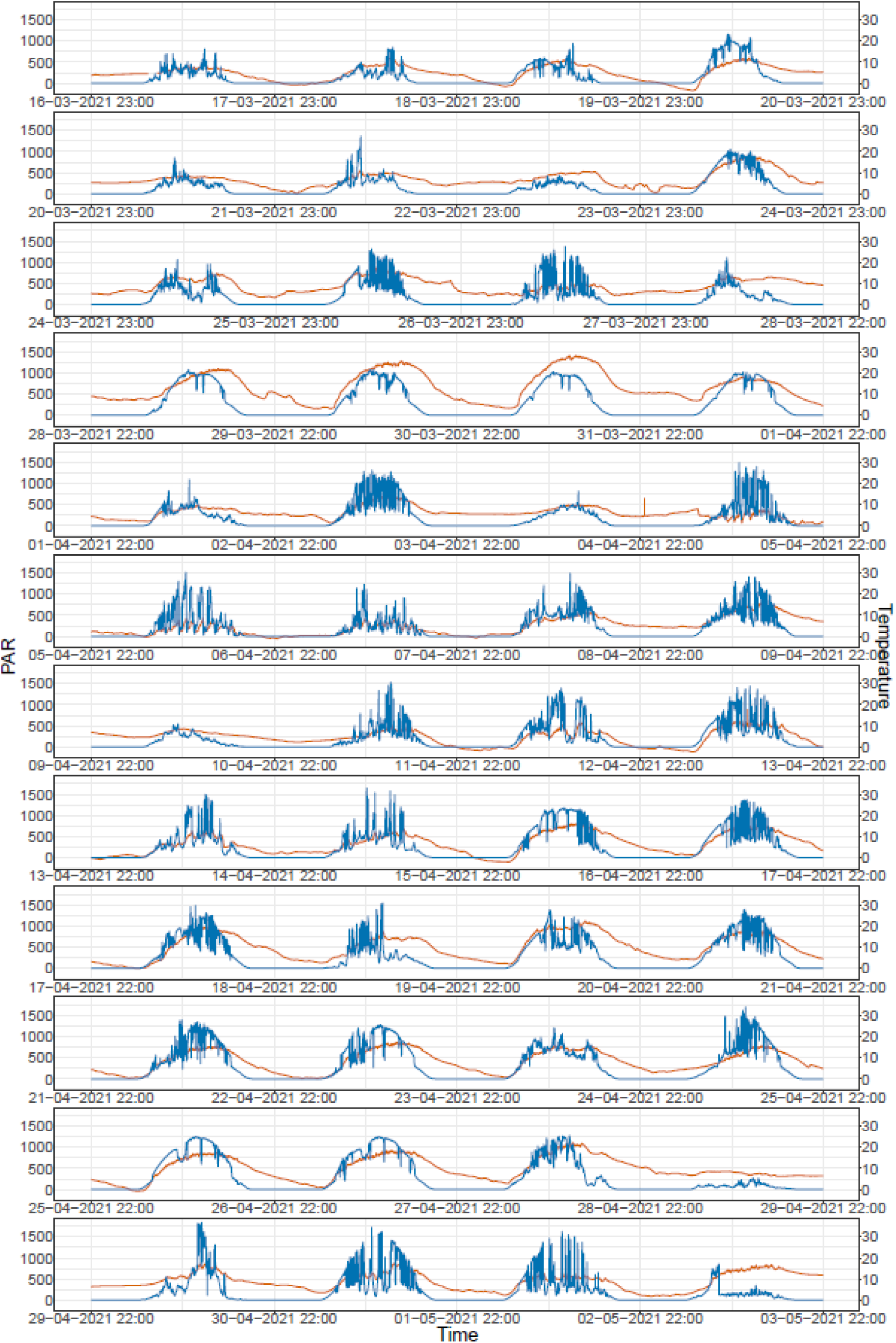
Light intensity and temperature for the 2021 semi-protected tunnel experiment.

## References

Alonso JM, Stepanova AN, Leisse TJ, et al. 2003. Genome-wide insertional mutagenesis of Arabidopsis thaliana. Science 301, 653–657.

Anderson JM, Chow WS, Park Y-I. 1995. The grand design of photosynthesis: Acclimation of the photosynthetic apparatus to environmental cues. Photosynthesis Research 46, 129–139.

Arends D, Arends D, Prins P, Broman KW, Jansen RC. 2014. Tutorial-multiple-QTL mapping (MQM) analysis Tutorial-Multiple-QTL Mapping (MQM) Analysis for R/qtl.

Bazakos C, Hanemian M, Trontin C, Jiménez-Gómez JM, Loudet O. 2017. New Strategies and Tools in Quantitative Genetics: How to Go from the Phenotype to the Genotype. Annual Review of Plant Biology 68, 435–455.

Bezouw RFHM van, Keurentjes JJB, Harbinson J, Aarts MGM. 2019. Converging phenomics and genomics to study natural variation in plant photosynthetic efficiency. The Plant Journal 97, 112–133.

Brachi B, Faure N, Horton M, Flahauw E, Vazquez A, Nordborg M, Bergelson J, Cuguen J, Roux F. 2010. Linkage and association mapping of Arabidopsis thaliana flowering time in nature. PLoS genetics 6, e1000940.

Broman KW, Wu H, Saunak Sen’, Churchill GA. 2003. R/qtl: QTL mapping in experimental crosses. BIOINFORMATICS APPLICATIONS NOTE 19, 889–890.

Chao M, Yin Z, Hao D, Zhang J, Song H, Ning A, Xu X, Yu D. 2014. Variation in Rubisco activase (RCAβ) gene promoters and expression in soybean [Glycine max (L.) Merr.]. Journal of Experimental Botany 65, 47–59.

Cruz JA, Savage LJ, Zegarac R, Hall CC, Satoh-Cruz M, Davis GA, Kovac WK, Chen J, Kramer DM. 2016. Dynamic Environmental Photosynthetic Imaging Reveals Emergent Phenotypes. Cell Systems 2, 365–377.

De Souza AP, Burgess SJ, Doran L, Hansen J, Manukyan L, Maryn N, Gotarkar D, Leonelli L, Niyogi KK, Long SP. 2022. Soybean photosynthesis and crop yield are improved by accelerating recovery from photoprotection. Science 377, 851–854.

Durand M, Matule B, Burgess AJ, Robson TM. 2021. Sunfleck properties from time series of fluctuating light. Agricultural and Forest Meteorology 308-309, 108554.

Durrett TP, Connolly EL, Rogers EE. 2006. Arabidopsis cpFtsY mutants exhibit pleiotropic defects including an inability to increase iron deficiency-inducible root Fe(III) chelate reductase activity. Plant Journal 47, 467–479.

El-Lithy ME, Rodrigues GC, van Rensen JJS, Snel JFH, Dassen HJHA, Koornneef M, Jansen MAK, Aarts MGM, Vreugdenhil D. 2005. Altered photosynthetic performance of a natural Arabidopsis accession is associated with atrazine resistance. Journal of Experimental Botany 56, 1625–1634.

Ferguson JN, Fernandes SB, Monier B, et al. 2021. Machine learning-enabled phenotyping for GWAS and TWAS of WUE traits in 869 field-grown sorghum accessions. Plant Physiology 187, 1481–1500.

Flood & Theeuwen, Schneeberger K, Keizer P, et al. 2020. Reciprocal cybrids reveal how organellar genomes affect plant phenotypes. Nature Plants 6, 13–21.

Flood PJ, van Heerwaarden J, Becker F, et al. 2016a. Whole-Genome Hitchhiking on an Organelle Mutation. Current Biology 26, 1306–1311.

Flood PJ, Kruijer W, Schnabel SK, et al. 2016b. Phenomics for photosynthesis, growth and reflectance in Arabidopsis thaliana reveals circadian and long-term fluctuations in heritability. Plant Methods 12, 14.

Forsberg SKG, Andreatta ME, Huang XY, Danku J, Salt DE, Carlborg Ö. 2015. The Multi-allelic Genetic Architecture of a Variance-Heterogeneity Locus for Molybdenum Concentration in Leaves Acts as a Source of Unexplained Additive Genetic Variance. PLoS Genetics 11, e1005648.

Fransz P, Linc G, Lee C-R, et al. 2016. Molecular, genetic and evolutionary analysis of a paracentric inversion in *Arabidopsis thaliana*. The Plant Journal 88, 159–178.

Furbank RT, Jimenez-Berni JA, George-Jaeggli B, Potgieter AB, Deery DM. 2019. Field crop phenomics: enabling breeding for radiation use efficiency and biomass in cereal crops. New Phytologist 223, 1714–1727.

Gaio D, Anantanawat K, To J, Liu M, Monahan L, Darling AEY. 2022. Hackflex: low-cost, high-throughput, Illumina Nextera Flex library construction. Microbial Genomics 8, 000744.

Garcia-Molina A, Leister D. 2020. Accelerated relaxation of photoprotection impairs biomass accumulation in Arabidopsis. Nature Plants 6, 9–12.

Garrison E, Marth G. 2012. Haplotype-based variant detection from short-read sequencing.

Giraut L, Falque M, Drouaud J, Pereira L, Martin OC, Mézard C. 2011. Genome-Wide Crossover Distribution in Arabidopsis thaliana Meiosis Reveals Sex-Specific Patterns along Chromosomes (M Lichten, Ed.). PLoS Genetics 7, e1002354.

Goto S, Mori H, Uchiyama K, Ishizuka W, Taneda H, Kono M, Kajiya-Kanegae H, Iwata H. 2021. Genetic Dissection of Growth and Eco-Physiological Traits Associated with Altitudinal Adaptation in Sakhalin Fir (Abies sachalinensis) Based on QTL Mapping. Genes 12, 1110.

He C, Holme J, Anthony J. 2014. SNP Genotyping: The KASP Assay. In: Fleury D,, In: Whitford R, eds. Methods in Molecular Biology. Crop Breeding: Methods and Protocols. New York, NY: Springer, 75–86.

Holt JS. 1990. Fitness and Ecological Adaptability of Herbicide-Resistant Biotypes. ACS Symposium Series. Managing Resistance to Agrochemicals. American Chemical Society, 419–429.

Johnson MP, Davison PA, Ruban AV, Horton P. 2008. The xanthophyll cycle pool size controls the kinetics of non-photochemical quenching in Arabidopsis thaliana. FEBS Letters 582, 262–266.

Joynson R, Molero G, Coombes B, Gardiner LJ, Rivera-Amado C, Piñera-Chávez FJ, Evans JR, Furbank RT, Reynolds MP, Hall A. 2021. Uncovering candidate genes involved in photosynthetic capacity using unexplored genetic variation in Spring Wheat. Plant Biotechnology Journal 19, 1537–1552.

Jung HS, Niyogi KK. 2009. Quantitative Genetic Analysis of Thermal Dissipation in Arabidopsis. Plant Physiology 150, 977–986.

Kaiser E, Morales A, Harbinson J. 2018. Fluctuating light takes crop photosynthesis on a rollercoaster ride. Plant Physiology 176, 977–989.

Korte A, Farlow A. 2013. The advantages and limitations of trait analysis with GWAS: a review. Plant Methods 2013 9:1 9, 1–9.

Kromdijk J, Głowacka K, Leonelli L, Gabilly ST, Iwai M, Niyogi KK, Long SP. 2016. Improving photosynthesis and crop productivity by accelerating recovery from photoprotection. Science 354.

Kromdijk J, Walter J. 2022. Relaxing non-photochemical quenching (NPQ) to improve photosynthesis in crops. Burleigh Dodds Science Publishing,.

Kruijer W, Boer MP, Malosetti M, Flood PJ, Engel B, Kooke R, Keurentjes JJB, van Eeuwijk FA. 2015. Marker-Based Estimation of Heritability in Immortal Populations. Genetics 199, 379–398.

Lee I, Bleecker A, Amasino R. 1993. Analysis of naturally occurring late flowering in Arabidopsis thaliana. Molecular and General Genetics MGG 237, 171–176.

Lehretz GG, Schneider A, Leister D, Sonnewald U. 2022. High non-photochemical quenching of VPZ transgenic potato plants limits CO _2_ assimilation under high light conditions and reduces tuber yield under fluctuating light. Journal of Integrative Plant Biology.

Levine RP. 1968. Genetic dissection of photosynthesis. Science 162, 768–771.

Li H. 2013. Aligning sequence reads, clone sequences and assembly contigs with BWA-MEM.

Li H, Handsaker B, Wysoker A, Fennell T, Ruan J, Homer N, Marth G, Abecasis G, Durbin R. 2009. The Sequence Alignment/Map format and SAMtools. Bioinformatics 25, 2078–2079.

Long SP, Taylor SH, Burgess SJ, Carmo-Silva E, Lawson T, Souza APD, Leonelli L, Wang Y. 2022. Into the Shadows and Back into Sunlight: Photosynthesis in Fluctuating Light. https://doi.org/10.1146/annurev-arplant-070221-024745 73, 617–648.

Martin M. 2011. Cutadapt removes adapter sequences from high-throughput sequencing reads. EMBnet.journal 17, 10–12.

McKenna A, Hanna M, Banks E, et al. 2010. The genome analysis toolkit: A MapReduce framework for analyzing next-generation DNA sequencing data. Genome Research 20, 1297–1303.

Meinke D, Sweeney C, Muralla R. 2009. Integrating the Genetic and Physical Maps of Arabidopsis thaliana: Identification of Mapped Alleles of Cloned Essential (EMB) Genes. PLOS ONE 4, e7386.

Morales F, Ancín M, Fakhet D, González-Torralba J, Gámez AL, Seminario A, Soba D, Ben Mariem S, Garriga M, Aranjuelo I. 2020. Photosynthetic Metabolism under Stressful Growth Conditions as a Bases for Crop Breeding and Yield Improvement. Plants 9, 88.

Müller P, Li X-P, Niyogi KK. 2001. Non-Photochemical Quenching. A Response to Excess Light Energy1. Plant Physiology 125, 1558–1566.

Murata N, Takahashi S, Nishiyama Y, Allakhverdiev SI. 2007. Photoinhibition of photosystem II under environmental stress. Biochimica et Biophysica Acta (BBA) - Bioenergetics 1767, 414–421.

Murchie EH, Kefauver S, Araus JL, Muller O, Rascher U, Flood PJ, Lawson T. 2018. Measuring the dynamic photosynthome. Annals of Botany 122, 207–220.

Oakley CG, Savage L, Lotz S, Larson GR, Thomashow MF, Kramer DM, Schemske DW. 2018a. Genetic basis of photosynthetic responses to cold in two locally adapted populations of Arabidopsis thaliana. Journal of Experimental Botany 69, 699–709.

Oakley CG, Savage L, Lotz S, Larson GR, Thomashow MF, Kramer DM, Schemske DW. 2018b. Genetic basis of photosynthetic responses to cold in two locally adapted populations of Arabidopsis thaliana. Journal of Experimental Botany 69, 699–709.

Oettmeier W. 1999. Herbicide resistance and supersensitivity in photosystem II. Cellular and Molecular Life Sciences CMLS 55, 1255–1277.

Ort DR, Merchant SS, Alric J, et al. 2015. Redesigning photosynthesis to sustainably meet global food and bioenergy demand. Proceedings of the National Academy of Sciences of the United States of America 112, 8529–8536.

Ortiz D, Hu J, Salas Fernandez MG. 2017. Genetic architecture of photosynthesis in Sorghum bicolor under non-stress and cold stress conditions. Journal of Experimental Botany 68, 4545–4557.

Poormohammad Kiani S, Maury P, Sarrafi A, Grieu P. 2008. QTL analysis of chlorophyll fluorescence parameters in sunflower (Helianthus annuus L.) under well-watered and water-stressed conditions. Plant Science 175, 565–573.

Prinzenberg AE, Campos-Dominguez L, Kruijer W, Harbinson J, Aarts MGM. 2020. Natural variation of photosynthetic efficiency in Arabidopsis thaliana accessions under low temperature conditions. Plant Cell and Environment 43, 2000–2013.

Ravi M, Chan SWL. 2010. Haploid plants produced by centromere-mediated genome elimination. Nature 464, 615–618.

Ravi M, Marimuthu MPA, Tan EH, et al. 2014. A haploid genetics toolbox for Arabidopsis thaliana. Nature communications 5, 5334.

Rochaix J-D. 2004. Genetics of the Biogenesis and Dynamics of the Photosynthetic Machinery in Eukaryotes. The Plant Cell 16, 1650.

Van Rooijen R, Kruijer W, Boesten R, Van Eeuwijk FA, Harbinson J, Aarts MGM. 2017. Natural variation of YELLOW SEEDLING1 affects photosynthetic acclimation of Arabidopsis thaliana. Nature Communications 8, 1–9.

Ruban AV. 2017. Crops on the fast track for light. Nature 541, 36–37.

Rungrat T, Almonte AA, Cheng R, Gollan PJ, Stuart T, Aro EM, Borevitz JO, Pogson B, Wilson PB. 2019. A Genome-Wide Association Study of Non-Photochemical Quenching in response to local seasonal climates in Arabidopsis thaliana. Plant Direct 3, e00138.

Rutherford AW, Osyczka A, Rappaport F. 2012. Back-reactions, short-circuits, leaks and other energy wasteful reactions in biological electron transfer: redox tuning to survive life in O(2). FEBS letters 586, 603–616.

Salomé PA, Bomblies K, Fitz J, Laitinen RAE, Warthmann N, Yant L, Weigel D. 2011. The recombination landscape in Arabidopsis thaliana F2 populations. Heredity 2012 108:4 108, 447–455.

Scheller HV, Jensen PE, Haldrup A, Lunde C, Knoetzel J. 2001. Role of subunits in eukaryotic Photosystem I. Biochimica et Biophysica Acta - Bioenergetics 1507, 41–60.

Schönfeld M, Yaacoby T, Michael O, Rubin B. 1987. Triazine Resistance without Reduced Vigor in Phalaris paradoxa. Plant Physiology 83, 329–333.

Shindo C, Aranzana MJ, Lister C, Baxter C, Nicholls C, Nordborg M, Dean C. 2005. Role of FRIGIDA and FLOWERING LOCUS C in Determining Variation in Flowering Time of Arabidopsis. Plant Physiology 138, 1163–1173.

Simon M, Loudet O, Durand S, Bérard A, Brunel D, Sennesal FX, Durand-Tardif M, Pelletier G, Camilleri C. 2008. Quantitative trait loci mapping in five new large recombinant inbred line populations of Arabidopsis thaliana genotyped with consensus single-nucleotide polymorphism markers. Genetics 178, 2253–2264.

Sloan DB, Wu Z, Sharbrough J. 2018. Correction of Persistent Errors in Arabidopsis Reference Mitochondrial Genomes. The Plant cell 30, 525–527.

Theeuwen TPJM, Lawson A, Tijink D, et al. 2022a. The NDH complex reveals a trade-off preventing maximizing photosynthesis in Arabidopsis thaliana.

Theeuwen TPJM, Logie LL, Harbinson J, Aarts MGM. 2022b. Genetics as a key to improving crop photosynthesis. Journal Of Experimental Botany.

Tietz S, Hall CC, Cruz JA, Kramer DM. 2017. NPQt: a chlorophyll fluorescence parameter for rapid estimation and imaging of non-photochemical quenching of excitons in photosystem-II-associated antenna complexes. Plant, Cell & Environment 40, 1243–1255.

Tzvetkova-Chevolleau T, Hutin C, Noël LD, et al. 2007. Canonical Signal Recognition Particle Components Can Be Bypassed for Posttranslational Protein Targeting in Chloroplasts. The Plant Cell 19, 1635–1648.

Visscher PM, Hill WG, Wray NR. 2008. Heritability in the genomics era — concepts and misconceptions. Nature Reviews Genetics 9, 255–266.

Walter B, Pieta T, Schünemann D. 2015. Arabidopsis thaliana mutants lacking cpFtsY or cpSRP54 exhibit different defects in photosystem II repair. Frontiers in Plant Science 6.

Wang Q, Zhao H, Jiang J, Xu J, Xie W, Fu X, Liu C, He Y, Wang G. 2017. Genetic architecture of natural variation in rice nonphotochemical quenching capacity revealed by genome-wide association study. Frontiers in Plant Science 8, 1773.

Warwick SI. 1991. Herbicide Resistance in Weedy Plants: Physiology and Population Biology. Annual Review of Ecology and Systematics 22, 95–114.

Wijnker E, Deurhof L, van de Belt J, et al. 2014. Hybrid recreation by reverse breeding in Arabidopsis thaliana. Nature protocols 9, 761–72.

Zhu XG, Long SP, Ort DR. 2010. Improving photosynthetic efficiency for greater yield. Annual Review of Plant Biology 61, 235–261.

Zhu XG, Ort DR, Whitmarsh J, Long SP. 2004. The slow reversibility of photosystem II thermal energy dissipation on transfer from high to low light may cause large losses in carbon gain by crop canopies: A theoretical analysis. Journal of Experimental Botany. Oxford Academic, 1167–1175.

